# c-di-AMP signaling is required for bile salts resistance and long-term colonization by *Clostridioides difficile*

**DOI:** 10.1101/2021.08.23.457418

**Authors:** Marine Oberkampf, Audrey Hamiot, Pamela Altamirano-Silva, Paula Bellés-Sancho, Yannick D. N. Tremblay, Nicholas DiBenedetto, Roland Seifert, Olga Soutourina, Lynn Bry, Bruno Dupuy, Johann Peltier

## Abstract

To cause disease, the important human enteropathogen *Clostridioides difficile* must colonize the gastro-intestinal tract but little is known on how this organism senses and responds to the harsh host environment to adapt and multiply. Nucleotide second messengers are signaling molecules used by bacteria to respond to changing environmental conditions. In this study, we showed for the first time that c-di-AMP is produced by *C. difficile* and controls the uptake of potassium, making it essential for growth. We found that c-di-AMP is involved in biofilm formation, cell wall homeostasis, osmotolerance as well as detergent and bile salt resistance in *C. difficile*. In a colonization mouse model, a strain lacking GdpP, a c-di-AMP degrading enzyme, failed to persist in the gut in contrast to the parental strain. We identified OpuR as a new regulator that binds c-di-AMP and represses the expression of the compatible solute transporter OpuC. Interestingly, an *opuR* mutant is highly resistant to a hyperosmotic or bile salt stress compared to the parental strain while an *opuCA* mutant is more susceptible A short exposure of *C. difficile* cells to bile salts resulted in a decrease of the c-di-AMP concentrations reinforcing the hypothesis that changes in membrane characteristics due to variations of the cellular turgor or membrane damages constitute a signal for the adjustment of the intracellular c-di-AMP concentration. Thus, c-di-AMP is a signaling molecule with pleiotropic effects that controls osmolyte uptake to confer osmotolerance and bile salt resistance in *C. difficile* and that is important for colonization of the host.

**One Sentence Summary:** c-di-AMP is an essential regulatory molecule conferring resistance to osmotic and bile salt stresses by controlling osmolyte uptake and contributing to gut persistence in the human enteropathogen *Clostridioides difficile*.

## INTRODUCTION

*Clostridioides difficile* is a medically important human enteropathogen that became a public health concern over the last two decades in industrialized countries (*1, 2*). This strict anaerobic spore-forming Gram-positive bacterium is a major cause of antibiotic-associated nosocomial diarrhoea in adults (*3*). Most virulent *C. difficile* strains produce two glycosylating toxins (TcdA and TcdB) which play a key role in disease pathogenesis by targeting the gut epithelium resulting in severe inflammation and damage to the colon (*4, 5*). Transmission of *C. difficile* is dependent on the production of highly resistant spores, which germinate in the small intestine in response to primary bile salts (*6, 7*). Normally the intestinal microbiota mediates colonization resistance against *C. difficile* but an antibiotic treatment disrupts the host microbiota, resulting in *C. difficile* growth, colonization of the intestine and toxin production (*8, 9*).

During the course of infection along the gastrointestinal tract of the host, *C. difficile* encounters multiple stresses, including numerous antimicrobial compounds, abrupt shifts in pH, reactive oxygen species produced during inflammation and the host immune response to infection (*10–12*). *C. difficile* vegetative cells are also exposed to primary and secondary bile salts. Primary bile salts produced by the human liver consists mainly s of cholate and chenodeoxycholate conjugated with either taurine or glycine. Secondary bile salts, predominantly comprising of deoxycholate and lithocholate in humans, are derived from primary bile salts by modifications carried out by intestinal bacteria (*13*). While the primary bile salt taurocholate induces spore germination, the secondary bile salt deoxycholate is a poor germinant and inhibits vegetative cell growth (*14*). Another important stress in the intestinal lumen is the high osmolarity (equivalent to 300 mM sodium chloride (NaCl)) (*15*).

Bacteria respond to osmotic stresses by adjusting their intracellular concentrations of osmolytes to limit transmembrane water fluxes and maintain turgor. The emergency response against an hyperosmotic shock is the uptake of potassium (K^+^) and it is followed by the synthesis and/or import of compatible solutes, such as carnitine and glycine betaine that act as osmoprotectants (*16*). These generally neutral compounds are preferred osmolytes because they can accumulate to very high concentrations without inducing severe disturbances in cellular metabolism (*17*). Several osmolyte transport systems have been identified in Gram-positive bacteria and interestingly, many of these transporters are controlled by the second messenger cyclic diadenosine monophosphate (c-di-AMP) (*18*). As an example, c-di-AMP inhibits the three K^+^ transport systems in *Bacillus subtilis* (the high affinities KimA and KtrAB and the low affinity KtrCD transport systems) (*19*), negatively controls the activity of the OpuC carnitine transporter in *Listeria monocytogenes* and *Staphylococcus aureus* or binds to the BusR repressor controlling the expression of the glycine betaine transporter genes *busAA-busAB* in *Lactococcus lactis* and *Streptococcus agalactiae* (*20, 21*).

C-di-AMP is widely produced among Gram-positive bacteria with many c-di-AMP synthesizing organisms being prominent human pathogens. C-di-AMP is synthesized from two molecules of ATP by di-adenylate cyclase (DAC) enzymes and degraded to pApA or AMP by distinct c-di-AMP phosphodiesterase (PDE) enzymes. Most important human pathogens possess only a single DAC domain containing protein called CdaA, which is essential for production of c-di-AMP. However, spore-forming Clostridia and Bacilli contain one (DisA) or two additional DACs (DisA and CdaS), respectively (*22*). DisA plays a role in the control of DNA integrity and CdaS is specifically involved in sporulation-related processes (*23–26*). Two other DACs CdaM and CdaZ are present only in few organisms (*27, 28*). Four different classes of PDEs degrade c-di-AMP but most of the reported PDEs belong to the membrane-bound GdpP protein family, which consists of a signal regulatory module linked to a GGDEF domain and a DHH-DHHA1 catalytic domain (*29*) (*30, 31*) (*32*). C-di-AMP is essential for growth under standard laboratory conditions in most of the Firmicutes (*18, 20, 33-36*). However, recent studies revealed that c-di-AMP becomes dispensable, if the bacteria are cultivated on specific minimal media (*19, 20, 35*). Moreover, intracellular accumulation of c-di-AMP is also toxic and inhibits growth (*37*). As a second messenger, c-di-AMP initiates signal transduction by binding to receptors that regulate downstream cellular processes. C-di-AMP receptors include enzymes such as the pyruvate carboxylase (PycA) in *L. monocytogenes* and *L. lactis* (*38, 39*), osmolyte transporters (*28, 40–46*), transcriptional regulators (*20, 21, 47*) and the KdpD sensor kinase of the KdpDE potassium-responsive two-component regulatory system, which regulates the expression of the KdpFABC K^+^ importer in *Staphylococcus aureus* (*48, 49*). C-di-AMP-responding riboswitches, formerly called *ydaO* riboswitches, have also been identified and control the expression of K^+^ uptake systems in *B. subtilis* and *Bacillus thuringiensis* (*19, 50, 51*). Besides the main and conserved function of c-di-AMP in osmoregulation, the second messenger has been implicated in a wide range of cellular processes including central metabolism, cell wall homeostasis, biofilm formation and virulence (*23, 30, 33, 36, 52-58*).

To understand how *C. difficile* interacts with its host, it is necessary to elucidate the survival mechanisms necessary for colonization and pathogenesis. In this study, weanalyzed c-di-AMP synthesis and degradation to show that c-di-AMP plays pleiotropic roles in *C. difficile* by controlling cell wall homeostasis, biofilm formation and persistence of the bacterium in the host intestine, as well as tolerance to osmotic, detergent and bile salts stresses. We also demonstrate that potassium homeostasis is an essential function regulated by c-di-AMP in *C. difficile*. We identify the c-di-AMP-regulated OpuR transcriptional repressor that connects osmotic and bile salt tolerance. We show that the OpuC encoding genes are the main target of BusR-mediated repression and that the OpuC transporter is a functional carnitine and glycine betaine uptake system. Furthermore, we report that bile salt exposure is rapidly sensed by the cells, resulting in a decrease of intracellular c-di-AMP concentrations. Together, these data suggest that c-di-AMP is a key regulatory molecule that modulates osmolyte uptake in response to osmotic and bile salts stresses sensing by the cells to regulate adaptation to the host intestinal environment during infection.

## RESULTS

### The second messenger c-di-AMP is produced in *C. difficile*

A bioinformatics search identified two genes encoding DACs (*CD0028* and *CD0110*), homologous to DisA and CdaA respectively, and one gene coding for a PDE (*CD3659*), homologous to GdpP in the genome of *C. difficile* 630. The presence of such enzymes strongly suggested that c-di-AMP might be produced by *C. difficile*. To test this hypothesis, nucleotides were extracted from *C. difficile* 630Δ*erm* grown to exponential phase and c-di-AMP was detected by LC-MS (Fig. 1A). In-frame deletion of the three genes encoding putative c-di-AMP turnover enzymes were then readily generated. In contrast, attempts to create a *disA*/*cdaA* double mutant strain in standard culture conditions were unsuccessful. This suggested that c-d-AMP might be essential for growth in rich medium in *C. difficile*, as demonstrated for other Gram-positive bacteria. To confirm the role of the predicted enzymes, the intracellular c-di-AMP concentrations of the different gene deletion strains were determined by LC-MS (Fig. 1A). Deletion of *gdpP* significantly enhanced the concentrations of c-di-AMP while the c-di-AMP concentrations were unchanged in a *disA* mutant and were reduced in a *cdaA* mutant, although the differences were not statistically significant.

**Figure 1:**
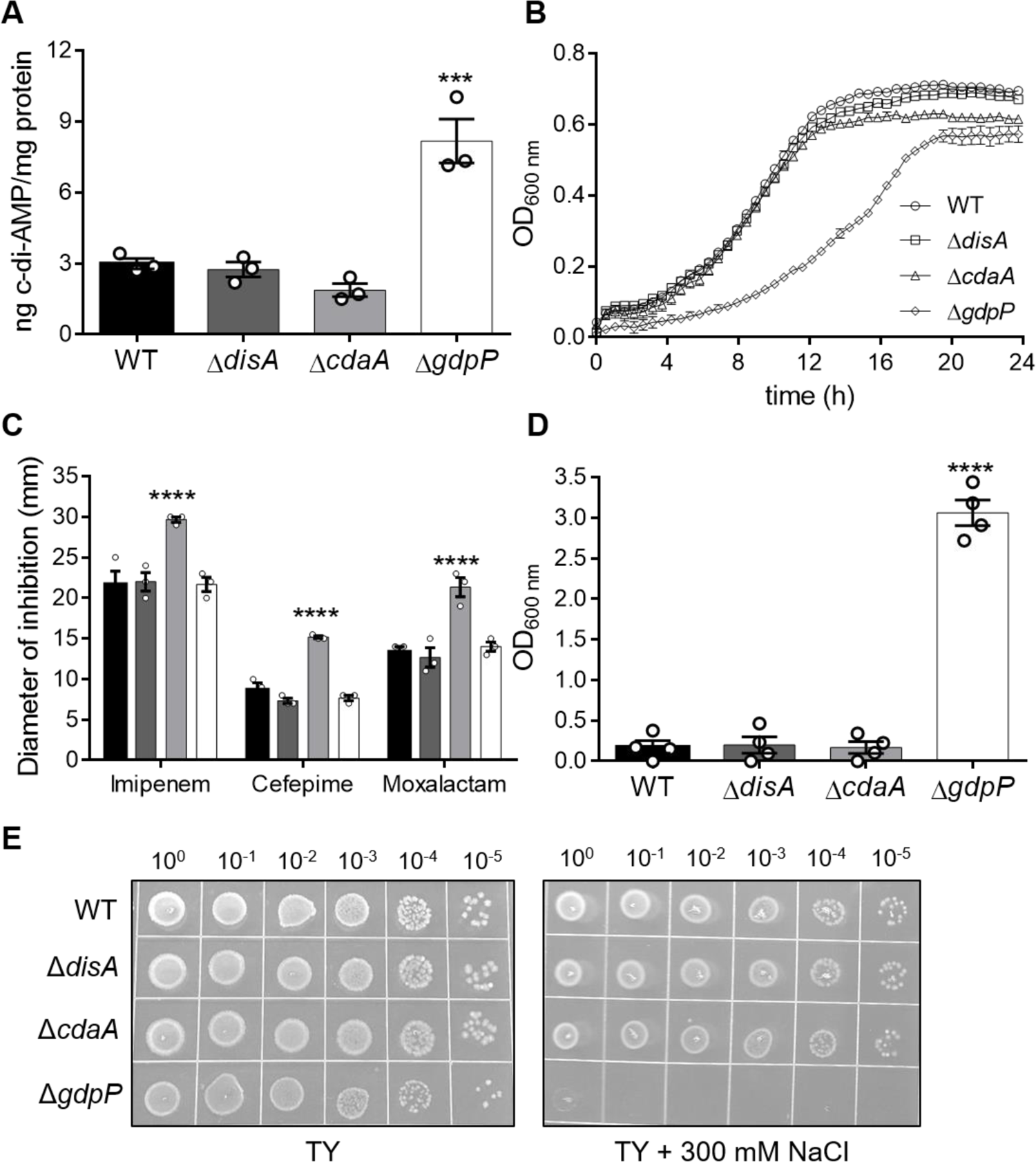
C-di-AMP concentrations in strains lacking c-di-AMP turnover enzymes and their associated changes in phenotypes. **(A)** Intracellular c-di-AMP concentrations in *C. difficile* wild type (WT), Δ*disA*, Δ*cdaA* and Δ*gdpP* strains grown in TY medium were quantified by LC-MS/MS. Means and standard error of the means (SEM) are shown; *n* = 3. *** *P* ≤ 0.001 by a one-way ANOVA followed by a Dunnett’s multiple comparison test comparing values to the average wild-type value. **(B)** Growth curves of *C. difficile* wild type (WT) and mutant strains in TY medium. Means and SEM are shown; *n* = 4. **(C)** Susceptibility of *C. difficile* wild type (black bars), Δ*disA* (dark grey bars), Δ*cdaA* (light grey bars) and Δ*gdpP* (white bars) strains to three β-lactam antibacterial drugs. Susceptibility was assessed using disk diffusion assays with 10 μg imipenem, 30 μg cefepime or 30 μg moxalactam. The zone of inhibition is expressed as the total diameter of the clearance zone and includes the diameter of filter paper disk (7 mm). Means and SEM are shown; *n* = 3. **** *P* ≤ 0.0001 by a two-way ANOVA followed by a Dunnett’s multiple comparison test comparing values to the average wild-type value. **(D)** Biofilm formation by *C. difficile* wild type and mutant strains grown in BHISG medium for 24 h. Means and SEM are shown; *n* = 4. **** *P* ≤ 0.0001 by a one-way ANOVA followed by a Dunnett’s multiple comparison test comparing values to the average wild-type value. **(E)** Growth of *C. difficile* wild type and mutant strains on TY agar or TY agar + 300 mM NaCl after 24 h incubation at 37°C. Spots (5μl) are from cells grown overnight in TY and serially diluted. Data are representative of 3 experiments.

### Fluctuations of c-di-AMP concentrations result in pleiotropic effects in *C. difficile*

To assess the role of c-di-AMP on the physiology of *C. difficile*, we investigated several phenotypes associated with fluctuations of c-di-AMP concentrations in other bacteria. We first explored the impact of *disA*, *cdaA* or *gdpP* deletions on *C. difficile* growth. While growths of the *disA* and the *cdaA* mutants in TY medium were similar to that of the wild-type strain, the *gdpP* deletion resulted in a growth defect (Fig. 1B). This is in agreement with the toxicity of intracellular accumulation of c-di-AMP reported in other bacteria (*37*). To determine if c-di-AMP affects cell wall homeostasis in *C. difficile*, we measured susceptibility to cell wall-targeting β-lactam antibacterial drugs using antibiotic disk diffusion assays. *C. difficile* Δ*cdaA* mutant was more susceptible to all tested antibacterial drugs whereas the Δ*disA* and Δ*gdpP* strains had the same susceptibility as the wild type strain (Fig. 1C). These data suggest a role for c-di-AMP in regulating the cell wall structure. We then assessed the ability of the mutants to form biofilms. Under our experimental conditions, no biofilm was observed in the wild type, the Δ*cdaA* or the Δ*disA* strains (Fig. 1D). In contrast, a strong biofilm was obtained with the Δ*gdpP* strain, revealing an association between c-di-AMP concentrations and biofilm formation in *C. difficile*.

Since regulation of osmotic balance is a conserved and major function of c-di-AMP in bacteria (*59*), we next tested the tolerance of *C. difficile* 630Δ*erm*, Δ*disA*, Δ*cdaA* and Δ*gdpP* strains to a high osmotic stress. The Δ*gdpP* mutant was highly susceptible to 300 mM NaCl when compared to the wild type strain (Fig. 1E). In contrast, Δ*disA* and Δ*cdaA* behaved like the wild type. Consistently, expression of the *cdaA* gene on a plasmid under control of the inducible *P_tet_* promoter resulted in an increased susceptibility of 630Δ*erm* to NaCl compared to a control strain carrying an empty vector (Fig. S1). Overxpression of *gdpP* on a plasmid had no impact on osmotolerance. These data demonstrate that c-di-AMP plays an important role in osmotolerance in *C. difficile*.

### c-di-AMP binds to proteins involved in K^+^ uptake

In bacteria, osmotic homeostasis is regulated by c-di-AMP concentrations via diverse K^+^ transport systems. *In-silico* analyses revealed that two potential K^+^ transport systems, KtrAB (*CD0696-CD0697*) and KdpABC (*CD1591-CD1593*), whose expression is regulated by the two-component regulatory system KdpDE (*CD1829-*CD1830), are present in the genome of *C. difficile* 630. In *S. aureus*, c-di-AMP inhibits expression of *kdpABC* and activity of KtrAB by binding to the universal stress protein (USP) domain of the histidine kinase KdpD and to the RCK_C domain of KtrA, respectively (*42, 48*). These domains are conserved in *C. difficile* homologues. To probe the interaction between c-di-AMP and the proteins KdpD and KtrA in *C. difficile*, we performed a <u>d</u>ifferential <u>ra</u>dial <u>c</u>apillary <u>a</u>ction of ligand <u>a</u>ssay (DRaCALA).

The corresponding genes were expressed as recombinant proteins in *Escherichia coli* and whole-cell extracts were incubated with [^32^P]-labelled c-di-AMP. Radiolabelled c-di-AMP interacted with both proteins indicating that they are c-di-AMP target proteins (Fig. 2A). In addition, an excess of unlabeled c-di-AMP, but not of other tested unlabeled nucleotides, outcompeted [^32^P]-c-di-AMP for binding with KdpD or KtrA, demonstrating the binding specificity of c-di-AMP to the proteins. Taken together, these results strongly suggest that c-di-AMP regulated proteins that controls K^+^ transport in *C. difficile*.

**Figure 2:**
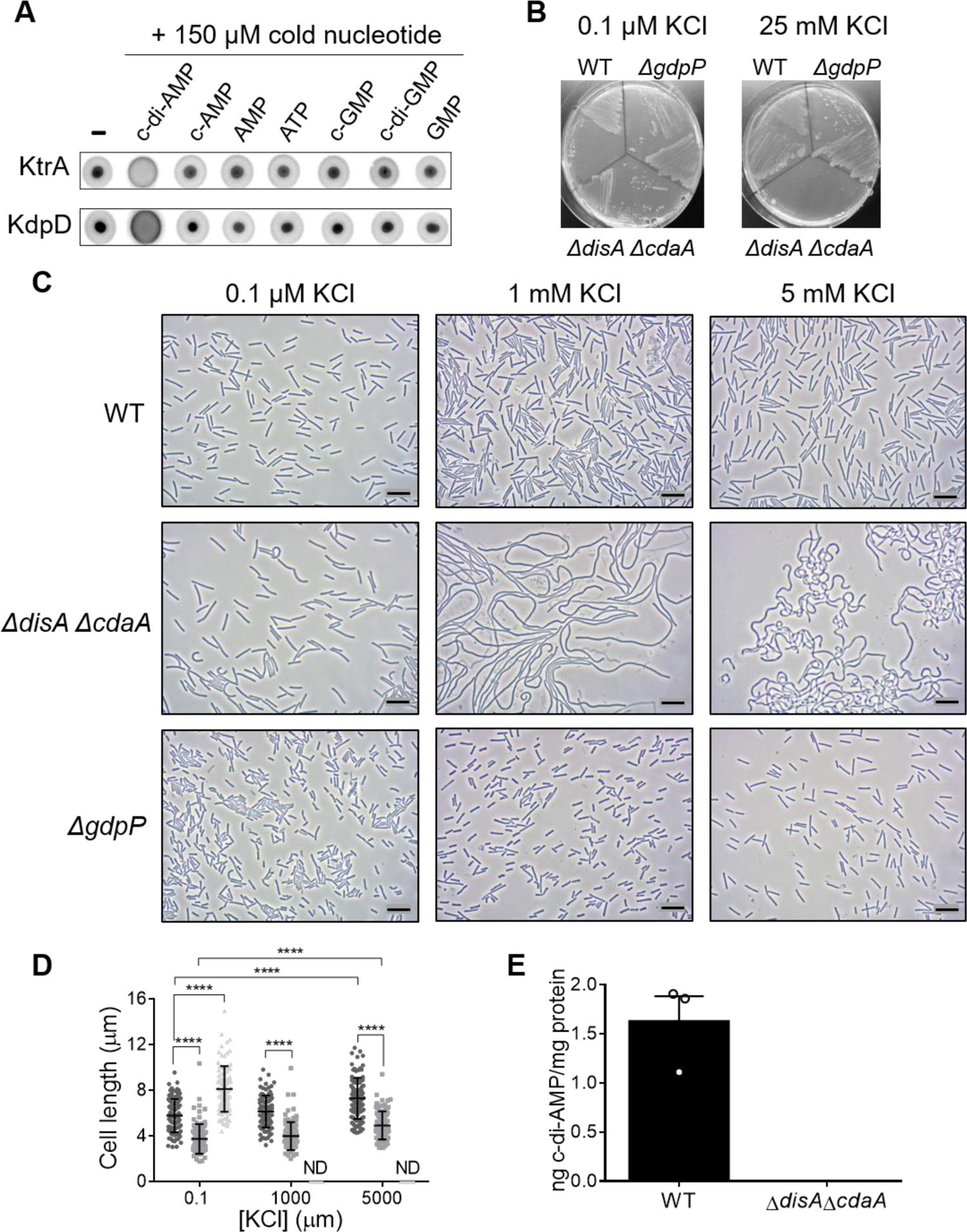
Control of K+ uptake is the essential function of c-di-AMP in *C. difficile*. **(A)** KtrA and KdpD bind [^32^P]-c-di-AMP in DRaCALAs. Binding of the radiolabeled ligand (1 nM) is indicated by dark spots centered on the nitrocellulose. In competition assays, excess of unlabeled nucleotides (150 μM) was added to the reaction before spotting on membrane. *n*=2 independent whole *E*. *coli* protein extracts. **(B)** Growth of *C. difficile* wild type, Δ*gdpP* and Δ*disA*Δ*cdaA* strains on CDMM agar plates containing 0.1 μM or 25 mM KCl after 48 h at 37 °C. Data are representative of 3 experiments. **(C)** Phase contrast microscopy image of *C. difficile* wild type, Δ*gdpP* and Δ*disA*Δ*cdaA* strains grown for 48 h at 37 °C on CDMM agar plates containing 0.1 μM, 1 mM or 5 mM KCl. Scale bars represent 10 μm. Data are representative of 3 experiments. **(D)** Scatter plots showing cell length with the median and standard deviation of each distribution indicated by a black line; *n* = 100. **** *P* ≤ 0.0001 by a two-way ANOVA followed by a Dunnett’s or Tukey’s multiple comparison test. ND= Not Determined. **(E)** Intracellular c-di-AMP concentrations in *C. difficile* wild type and Δ*disA*Δ*cdaA* grown on CDMM agar + 0.1 mM KCl were quantified by LC-MS/MS. Means and SEM are shown; *n* = 3.

### Regulation of K^+^ uptake is the essential function of c-di-AMP in *C. difficile*

Assuming that K^+^ homeostasis might be the essential function of c-di-AMP in *C. difficile*, we attempted to generate a Δ*disA*Δ*cdaA* double mutant by employing our gene deletion protocol using a modified *C. difficile* minimal medium (CDMM) containing 0.1 mM of K^+^ instead of BHI. The Δ*disA*Δ*cdaA* mutant was successfully generated and was viable under these conditions. However, growth of the mutant on plates was severely affected in presence of 5 mM of K^+^ and abolished in presence of 25 mM of K^+^ (Fig. 2B and S2A). Examination of Δ*disA*Δ*cdaA* cells by phase contrast microscopy also revealed a strong impact of the K^+^ concentration on cell morphology characterized by a pronounced elongation and curvature of the cells when K^+^ concentration increased (Fig. 2C and 2D). Importantly, c-di-AMP could not be detected by LC-MS in nucleotide extracts from this strain, indicating that DisA and CdaA are the only two enzymes involved in the production of c-di-AMP in *C. difficile* (Fig. 2E). In contrast to Δ*disA*Δ*cdaA*, the wild type and the Δ*gdpP* strains grew on CDMM plates with all K^+^ concentrations but their cell morphology was also affected by the K^+^ concentration (Fig. 2B and 2C). Cells of the wild type strain became shorter as K^+^ concentration decreased and cells of Δ*gdpP* were consistently shorter than those of wild type (Fig. 2C and 2D). In addition, Δ*gdpP* cell shape aberrations such as bent cells and cells with division defects were observed in presence of low K^+^ concentrations (Fig. S2B). All together, these data demonstrate that control of K^+^ uptake needs to be tightly regulated and is an essential function of c-di-AMP in *C. difficile*.

### The transcriptional regulator OpuR binds c-di-AMP and represses the expression of the osmolyte transporter OpuCAC encoding genes in *C. difficile*

In response to an hyperosmotic stress, the import of K^+^ into bacterial cells is followed by a secondary response involving the synthesis or the uptake of compatible solutes (*16*). Interestingly, *C. difficile* possesses an ortholog to the c-di-AMP protein BusR, a transcriptional repressor of the glycine betaine uptake system BusAB in *L. lactis* and *S. agalactiae* (*20, 21*). Using DRaCALA, we demonstrated the interaction between c-di-AMP and BusR-like of *C. difficile* (Fig. 3A). We then used RNA-seq to compare the transcriptomes of the Δ*gdpP* and the Δ*gdpP* Δ*busR*-like mutant strains. Using a fold change cutoff of >2-fold and a *P* value limit of 0.05, we identified 48 genes whose expression was dependent upon BusR-like, with 16 positively and 32 negatively regulated genes (Table 1). Of these genes, the most highly induced were two genes, *CD0900-0901*, comprising an apparent operon. These genes encode a putative compatible solute ABC transporter system composed of an ATP binding protein (*CD0900*, *opuCA*) and a permease (*CD0901*, *opuCC*) presenting a limited homology with BusAA (35.1% identity, 55,1% similarity) and BusAB (21.7% identity, 38,1% similarity) of *L. lactis*, respectively. The results of the RNA-seq analyses for *opuCAC* expression were confirmed by qRT-PCR (Fig. S3), demonstrating that the c-di-AMP binding protein BusR-like, hereafter named OpuR, is a repressor of the *opuCAC* operon in *C. difficile*.

**Figure 3:**
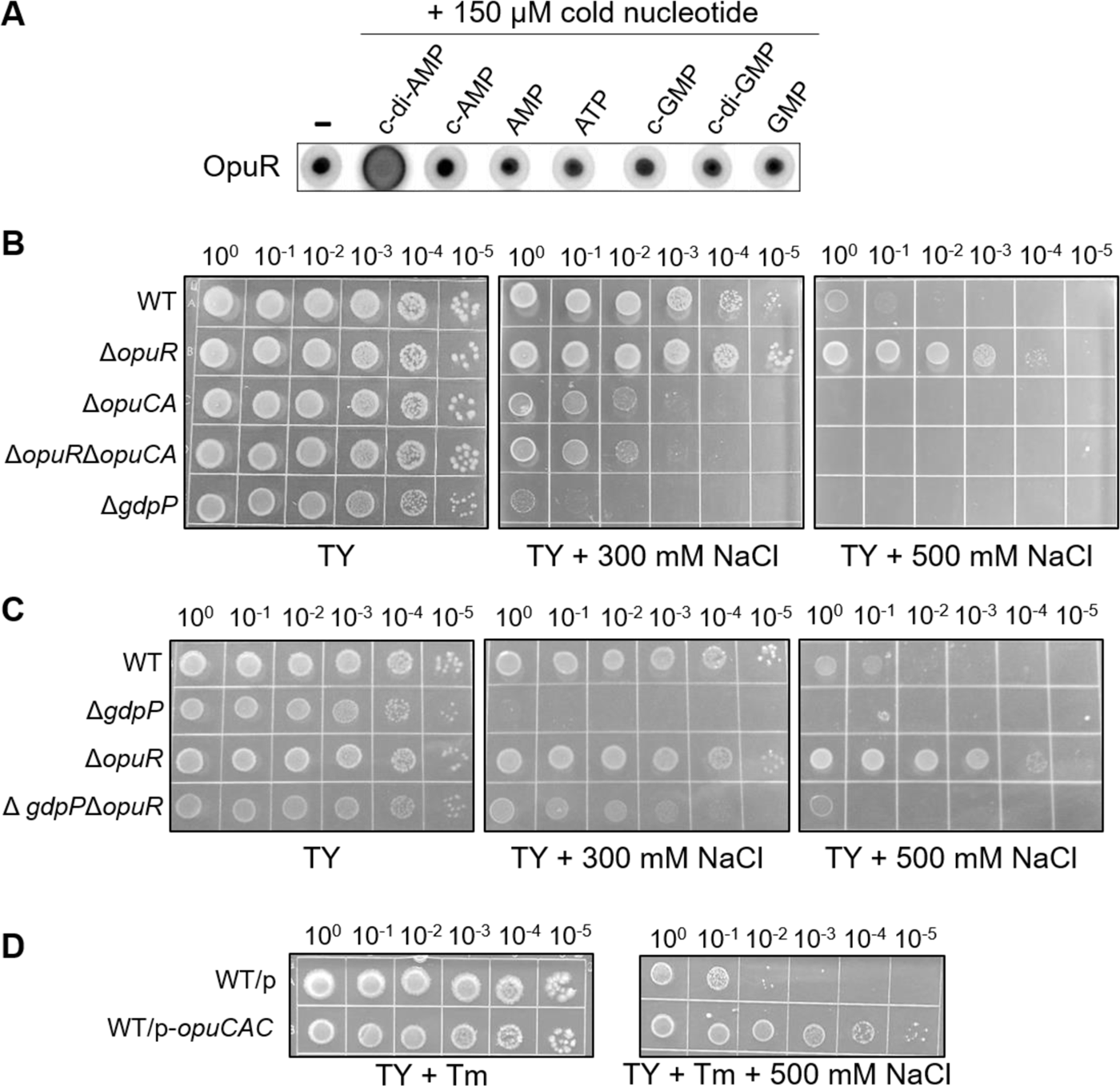
OpuR is a c-di-AMP target controlling osmotic homeostasis through repression of *opuCAC* expression. **(A)** OpuR binds [^32^P]-c-di-AMP in DRaCALAs. Binding of the radiolabeled ligand (1 nM) is indicated by dark spots centered on the nitrocellulose. In competition assays, excess of unlabeled nucleotides (150 μM) was added to the reaction before spotting on membrane. *n*=2 independent whole *E*. *coli* protein extracts. **(B)** and **(C)** Growth of B. *difficile* wild type and mutant strains on TY agar, TY agar + 300 mM NaCl or TY agar + 500 mM NaCl after 24 h incubation at 37°C. Spots (5μl) are from cells grown overnight in TY and serially diluted. Data are representative of 3 experiments. **(D)** Growth of *C. difficile* wild type strains carrying *p* (vector control) or *p*-*opuCAC* (constitutively expresses *opuCAC*) on TY agar + Tm or TY agar + Tm + 500 mM NaCl after 48 h incubation at 37°C. Spots (5μl) are from cells grown overnight in TY and serially diluted. Data are representative of 3 experiments.

**Table 1.**
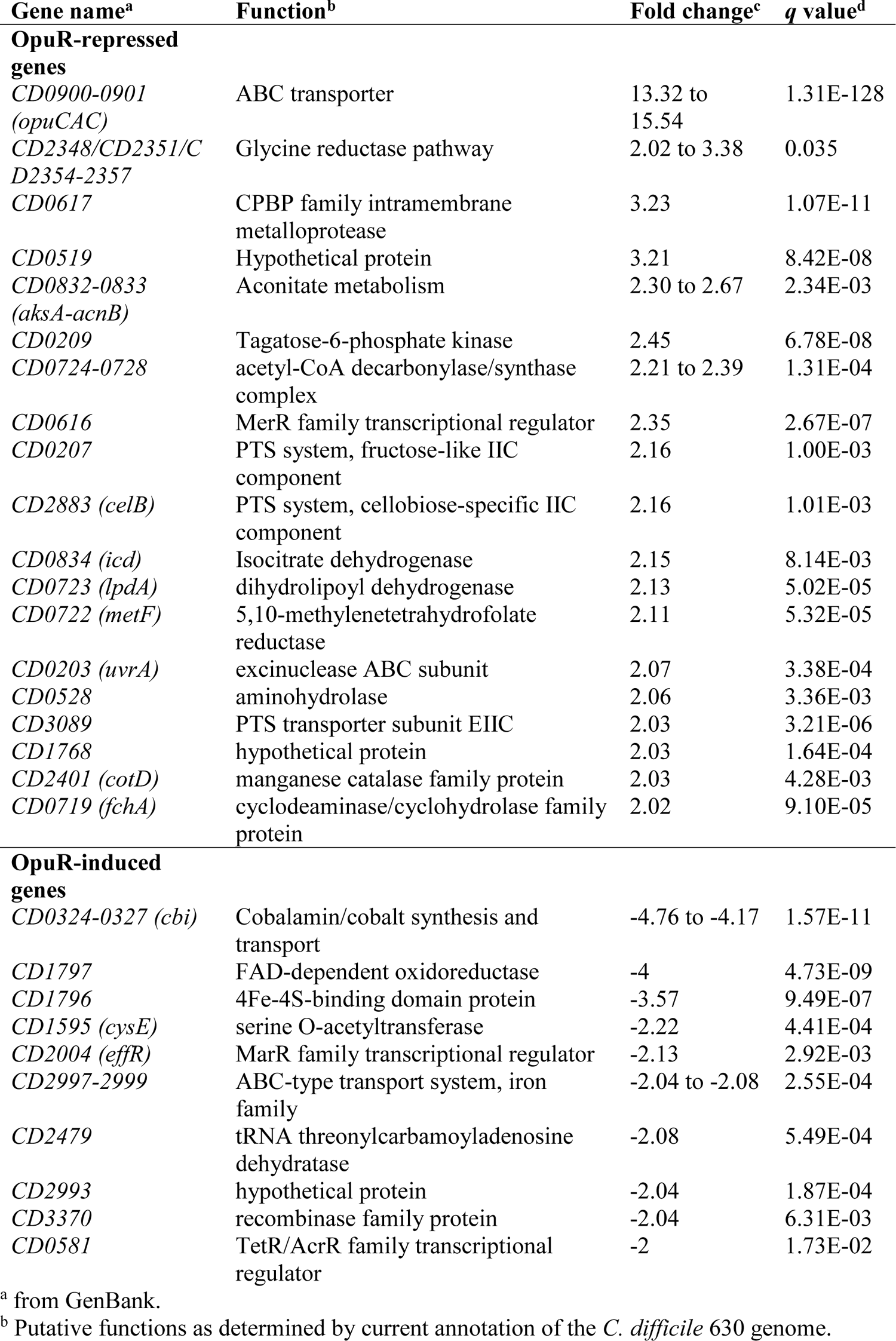
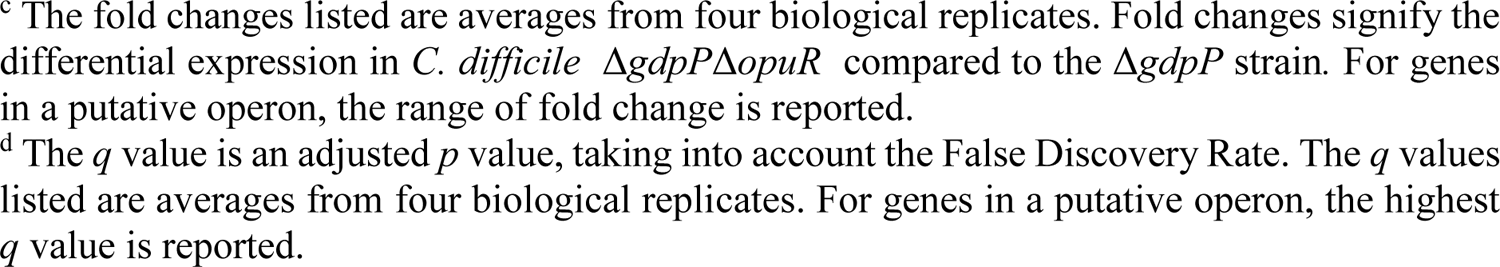
Genes regulated in the Δ*gdpP*Δ*opuR* strain in comparison to the Δ*gdpP* strain.

| Gene name<sup>a</sup> | Function<sup>b</sup> | Fold change<sup>c</sup> | q value<sup>d</sup> |
| --- | --- | --- | --- |
| **OpuR-repressed genes** |  |  |  |
| *CD0900-0901*<br>(*opuCAC*) | ABC transporter | 13.32 to 15.54 | 1.31E-128 |
| *CD2348/CD2351/CD2354-2357* | Glycine reductase pathway | 2.02 to 3.38 | 0.035 |
| *CD0617* | CPBP family intramembrane metalloprotease | 3.23 | 1.07E-11 |
| *CD0519* | Hypothetical protein | 3.21 | 8.42E-08 |
| *CD0832-0833*<br>(*aksA-acnB*) | Aconitate metabolism | 2.30 to 2.67 | 2.34E-03 |
| *CD0209* | Tagatose-6-phosphate kinase | 2.45 | 6.78E-08 |
| *CD0724-0728* | acetyl-CoA decarbonylase/synthase complex | 2.21 to 2.39 | 1.31E-04 |
| *CD0616* | MerR family transcriptional regulator | 2.35 | 2.67E-07 |
| *CD0207* | PTS system, fructose-like IIC component | 2.16 | 1.00E-03 |
| *CD2883 (celB)* | PTS system, cellobiose-specific IIC component | 2.16 | 1.01E-03 |
| *CD0834 (icd)* | Isocitrate dehydrogenase | 2.15 | 8.14E-03 |
| *CD0723 (lpdA)* | dihydrolipoyl dehydrogenase | 2.13 | 5.02E-05 |
| *CD0722 (metF)* | 5,10-methylenetetrahydrofolate reductase | 2.11 | 5.32E-05 |
| *CD0203 (uvrA)* | excinuclease ABC subunit | 2.07 | 3.38E-04 |
| *CD0528* | aminohydrolase | 2.06 | 3.36E-03 |
| *CD3089* | PTS transporter subunit EIIC | 2.03 | 3.21E-06 |
| *CD1768* | hypothetical protein | 2.03 | 1.64E-04 |
| *CD2401 (cotD)* | manganese catalase family protein | 2.03 | 4.28E-03 |
| *CD0719 (fchA)* | cyclodeaminase/cyclohydrolase family protein | 2.02 | 9.10E-05 |
| **OpuR-induced genes** |  |  |  |
| *CD0324-0327 (cbi)* | Cobalamin/cobalt synthesis and transport | -4.76 to -4.17 | 1.57E-11 |
| *CD1797* | FAD-dependent oxidoreductase | -4 | 4.73E-09 |
| *CD1796* | 4Fe-4S-binding domain protein | -3.57 | 9.49E-07 |
| *CD1595 (cysE)* | serine O-acetyltransferase | -2.22 | 4.41E-04 |
| *CD2004 (effR)* | MarR family transcriptional regulator | -2.13 | 2.92E-03 |
| *CD2997-2999* | ABC-type transport system, iron family | -2.04 to -2.08 | 2.55E-04 |
| *CD2479* | tRNA threonylcarbamoyladenosine dehydratase | -2.08 | 5.49E-04 |
| *CD2993* | hypothetical protein | -2.04 | 1.87E-04 |
| *CD3370* | recombinase family protein | -2.04 | 6.31E-03 |
| *CD0581* | TetR/AcrR family transcriptional regulator | -2 | 1.73E-02 |

<sup>a</sup> from GenBank.
<sup>b</sup> Putative functions as determined by current annotation of the *C. difficile* 630 genome.
<sup>c</sup> The fold changes listed are averages from four biological replicates. Fold changes signify the differential expression in *C. difficile* $\Delta gdpP\Delta opuR$ compared to the $\Delta gdpP$ strain. For genes in a putative operon, the range of fold change is reported.
<sup>d</sup> The *q* value is an adjusted *p* value, taking into account the False Discovery Rate. The *q* values listed are averages from four biological replicates. For genes in a putative operon, the highest *q* value is reported.

### OpuR and OpuCAC play an important role in osmotic homeostasis in *C. difficile*

To evaluate the role played by OpuR and OpuCAC in osmotolerance, we analyzed the ability of Δ*opuCA* and Δ*opuR* to grow in the presence of NaCl. While Δ*opuCA* was more susceptible to a hyperosmotic stress than the wild type strain but less than Δ*gdpP*, deletion of *opuR* resulted in a strong increase in resistance to 500 mM NaCl (Fig. 3B). In addition, a Δ*opuR*Δ*opuCA* double mutant was as susceptible to NaCl as the *opuCA* single mutant strain and deletion of *opuR* in the Δ*gdpP* background partially rescued osmoresistance (Fig. 3B and 3C). This indicates that the osmoresistance phenotype of Δ*opuR* is fully mediated by the derepression of *opuCAC* genes expression and that the osmosusceptibility of Δ*gdpP* is not only caused by the inhibition of *opuC* expression. Then, to confirm that higher expression levels of OpuCAC were sufficient to increase osmoresistance, the vector p-*opuCAC*, expressing the *opuCAC* genes under the control of the strong and constitutive *P_cwp2_* promoter (*60*), was constructed and introduced into the wild-type strain. As expected, *C. difficile* overexpressing *opuCA*C was more resistant to an osmotic stress compared to the vector control strain (Fig. 3D). Taken together, these data suggest that the deletion of *opuR* results in increased osmolyte uptake through the derepression of *opuCAC* rendering the strain more resistant to an osmotic stress and indicate that the control of OpuR by c-di-AMP plays an important role in osmoresistance.

### OpuCAC is an osmolyte transporter

To provide evidence on the function of OpuCAC, we tested the growth of our deletion mutant panel on CDMM agar plates supplemented with 200 mM NaCl. Under these conditions, the wild type, Δ*opuR*, Δ*opuCA* and Δ*opuR*Δ*opuCA* were equally resistant to the hyperosmotic stress (Fig. 4A), suggesting that the osmoprotective compound transported by OpuCAC was not present in the medium. In contrast, Δ*gdpP* and Δ*gdpP*Δ*opuR* were more susceptible than the wild type, consistent with the presence of K^+^ in the medium. Interestingly, addition of 0.4 mM of glycine betaine or carnitine in the culture medium improved the growth of the wild type and the *opuR* mutant in the presence of NaCl but had no effect on the growth of Δ*opuCA* and Δ*opuR*Δ*opuCA* (Fig. 4A). Osmotolerance of Δ*gdpP*Δ*opuR* was also increased when compared to Δ*gdpP.* These results demonstrate that the phenotypes of Δ*opuR* and Δ*opuCA* in response to osmotic stress result from a dysregulated compatible solute uptake and that the OpuCAC transporter can take up both betaine and carnitine.

**Figure 4:**
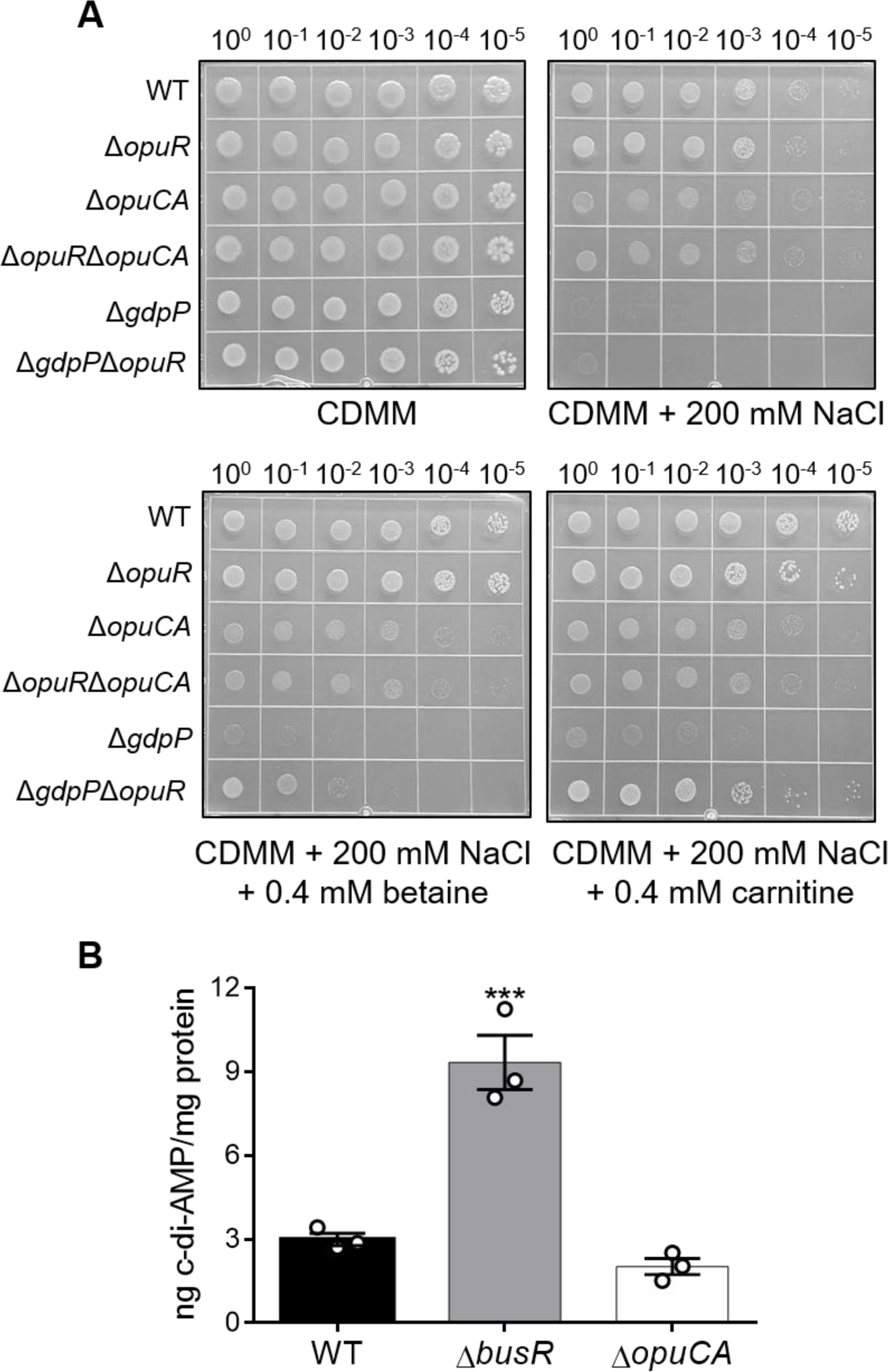
OpuCAC is an osmolyte transporter. **(A)** Growth of *C. difficile* wild type and mutant strains on CDMM agar, CDMM agar + 200 mM NaCl, CDMM agar + 200 mM NaCl + 0.4 mM glycine betaine or CDMM agar + 200 mM NaCl + 0.4 mM carnitine after 48 h incubation at 37°C. Spots (5μl) are from log-phase cells grown in TY, washed once in 0.9% saline and serially diluted. Data are representative of 3 experiments. **(B)** Intracellular c-di-AMP concentrations in *C. difficile* wild type, Δ*opuR* and Δ*opuCA* strains grown in TY medium were quantified by LC-MS/MS. Means and standard error of the means (SEM) are shown; *n* = 3. *** *P* ≤ 0.001 by a one-way ANOVA followed by a Dunnett’s multiple comparison test comparing values to the average wild-type value.

Next, we sought to determine if changes in compatible solute uptake caused by loss of OpuR or OpuCA affect cellular c-di-AMP concentrations (Fig. 4B). Measurement of intracellular c-di-AMP concentrations by LC-MS/MS revealed increased amounts of c-di-AMP in Δ*opuR* grown in TY compared with the wild type strain. In contrast, c-di-AMP concentrations were decreased in Δ*opuCA*, though the difference did not reach statistical significance. This suggests that c-di-AMP concentrations are modulated by the intracellular concentration of compatible solutes in *C. difficile*. Thus, an increased uptake of glycine betaine or carnitine leads to an elevation of the intracellular c-di-AMP concentration which will positively impact OpuR-mediated repression of the OpuCAC osmolyte transporter genes.

### OpuR mediates resistance to the detergent activity of bile salts

Several osmolyte uptake systems, including OpuC are involved in bile tolerance in *Listeria monocytogenes* (*61*). These data prompted us to investigate the role of OpuR and OpuCA in bile salt tolerance in *C. difficile*. We first analyzed the phenotypes of Δ*opuR*, Δ*opuCA* and Δ*opuR*Δ*opuCA* in response to a commercial bile salt extract. Surprisingly, Δ*opuR* was extremely resistant to this stress compared to the wild type strain, while Δ*opuCA* and Δ*opuR*Δ*opuCA* were more susceptible than the parental strain (Fig. 5A and S4A). Because bile salt extract is a mixture of primary and secondary bile salts, we then tested the response of the mutant strains to the primary bile salt cholate as well as the secondary bile acid deoxycholate, a cholate derivative. Results were similar for both bile salts and in line with those obtained with the bile salt extract; *opuR* deletion increased resistance to the two bile salts while *opuCA* deletion in the wild type or the Δ*opuR* background resulted in a hypersensitivity to these bile salts (Fig. 5A). Because bile salts have potent detergent properties, we tested the tolerance of the mutant strains to the ionic detergent SDS and the non-ionic detergent Triton X-100 (Fig. 5B). Again, Δ*opuR* was found to be highly resistant to both compounds, whereas Δ*opuCA* and Δ*opuR*Δ*opuCA* were more sensitive than the wild type strain.

**Figure 5:**
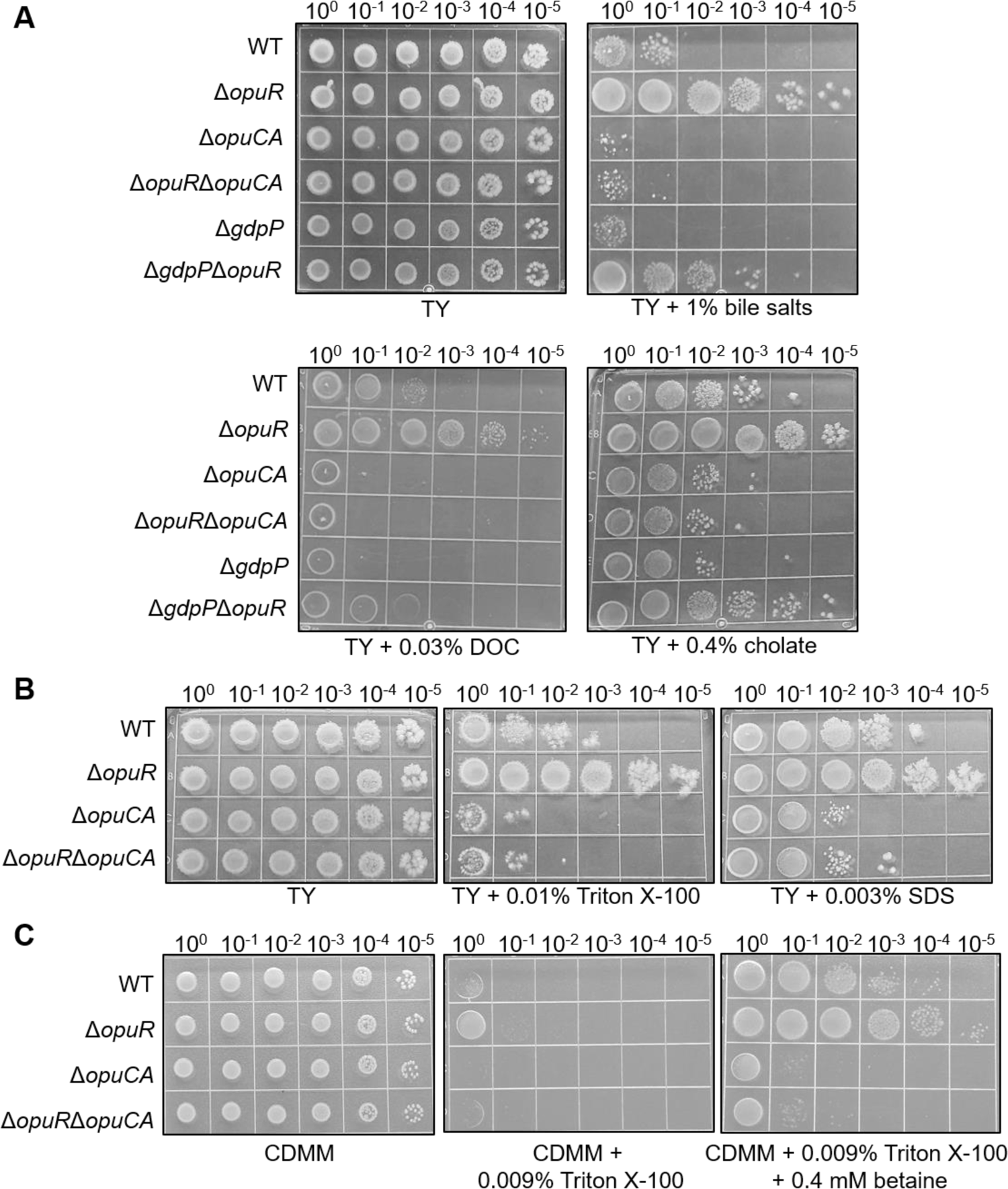
C-di-AMP modulates resistance to the detergent action of bile salts by controlling the OpuCAC-mediated import of osmolytes through OpuR. **(A)** Growth of *C. difficile* wild type and mutant strains on TY agar, TY agar + 1% bile salts, TY agar + 0.03% deoxycholate (DOC) or TY agar + 0.4% cholate after 48 h incubation at 37°C. Spots (5μl) are from cells grown overnight in TY and serially diluted. Data are representative of 3 experiments. **(B)** Growth of *C. difficile* wild type and mutant strains on TY agar, TY agar + 0.01% Triton X-100 or TY agar + 0.003% SDS after 48 h incubation at 37°C. Spots (5μl) are from cells grown overnight in TY and serially diluted. Data are representative of 3 experiments. **(C)** Growth of *C. difficile* wild type and mutant strains on CDMM agar, CDMM agar + 0.009% Triton X-100 or CDMM agar + 0.009% Triton X-100 + 0.4 mM glycine betaine after 48 h incubation at 37°C. Spots (5μl) are from log-phase cells grown in TY, washed once in 0.9% saline and serially diluted. Data are representative of 3 experiments.

We then sought to determine whether the import of compatible solutes into the cells by OpuCAC directly drove detergent tolerance. Wild type, Δ*opuR*, Δ*opuCA* and Δ*opuR*Δ*opuCA* mutants were grown on CDMM agar plates supplemented with 0.008 to 0.01 % Triton X-100 in absence or presence of 0.4 mM glycine betaine (Fig. 5C and S4B). Growth defect caused by the detergent was in the same range for all strains in absence of compatible solutes. Presence of glycine betaine in the medium partially restored the growth of the wild type and to a better extent the growth of Δ*opuR*. Strains Δ*opuCA* and Δ*opuR*Δ*opuCA* remained more susceptible to Triton X-100 than the wild type strain but their growth was also slightly improved by glycine betaine, suggesting another osmolyte transporter is present. Taken together, these results show that OpuR plays an important role in resistance to the detergent activity of bile salts by controlling the transport of compatible solutes by OpuCAC into *C. difficile* cells.

### C-di-AMP concentrations are reduced by bile salt exposure and modulate resistance to bile salts

To establish a functional link between c-di-AMP and bile salt resistance, phenotypes of Δ*gdpP* and Δ*gdpP*Δ*opuR* in response to either a bile salt extract, cholate or deoxycholate were also investigated (Fig. 5A). For all tested bile salts, Δ*gdpP* showed the same susceptibility phenotype as Δ*opuCA* and deletion of *opuR* in the Δ*gdpP* mutant abolished the susceptibility. These data indicate that c-di-AMP modulates bile salt resistance primarily through the repressing activity of OpuR on *opuCAC* expression.

We next examined the effect of bile salts on c-di-AMP concentrations in 630Δ*erm*. Cells grown to the exponential phase were washed and resuspended in a low osmolarity buffer with glucose.

After a 10 min incubation, water, NaCl, deoxycholate or cholate was added and cells were incubated another 10 min. C-di-AMP was then quantified in the different samples using a competitive ELISA assay (Fig. 6). In our conditions, exposure to NaCl resulted in a slight decrease of the c-di-AMP concentrations in comparison to the water-control. In contrast, a rapid and statistically significant degradation of c-di-AMP was observed in the samples treated with deoxycholate and cholate relative to the water control. This demonstrates that bile salt exposure promotes c-di-AMP degradation in *C. difficile*.

**Figure 6:**
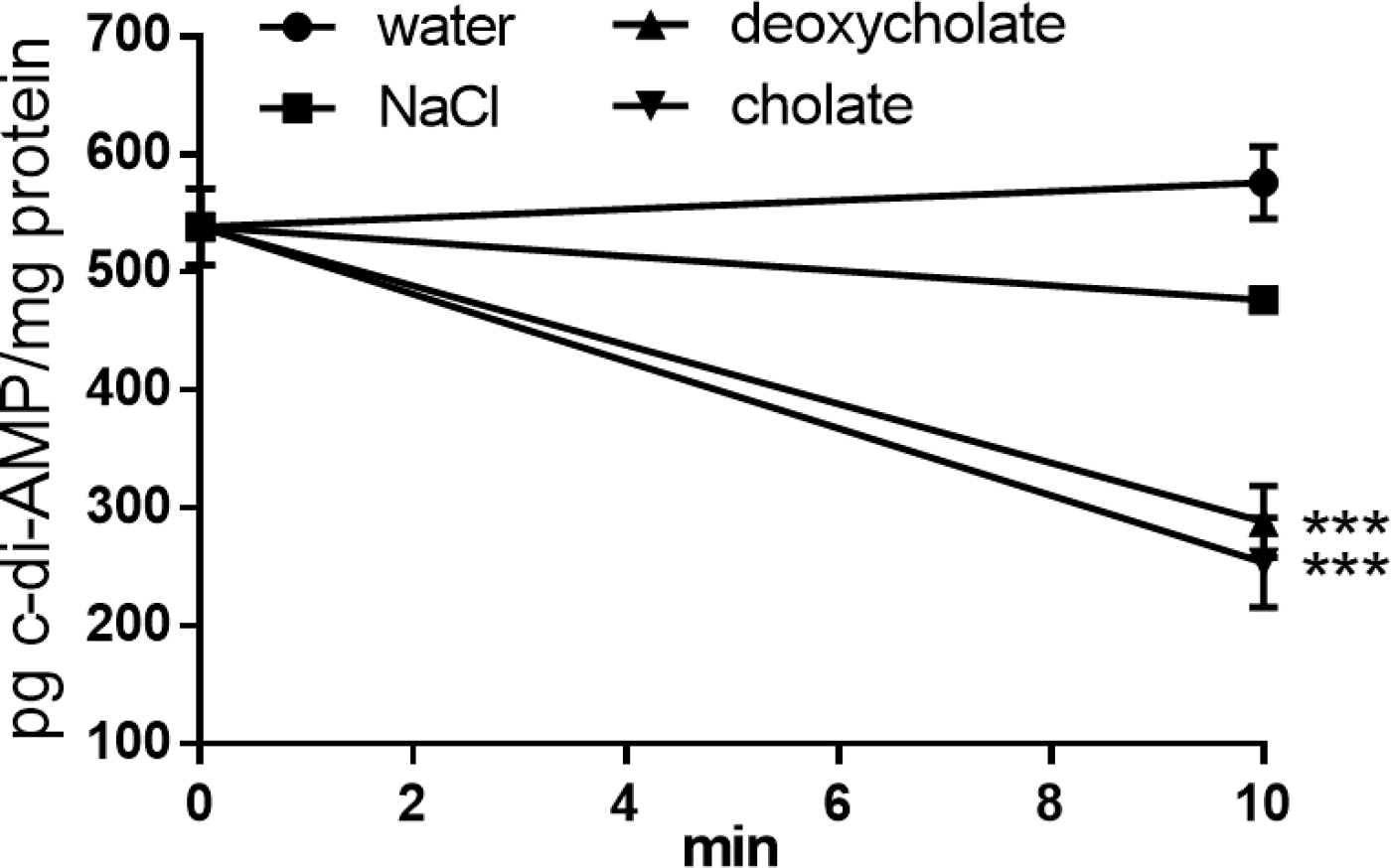
A bile salt treatment decreases c-di-AMP concentrations in *C. difficile*. *C. difficile* Δ*erm* cells suspended and incubated for 10 min in a low osmolarity buffer with glucose (t_0_) were treated with water, 300 mM NaCl, 0.028% deoxycholate or 0.4% cholate for 10 min at 37°C (t_10_). c-di-AMP concentrations were measured at t_0_ and t_10_ using a competitive c-di-AMP ELISA. Means and standard error of the means (SEM) from 3 biological replicates were plotted as pg c-di-AMP/ mg protein. *** *P* ≤ 0.001 by a one-way ANOVA followed by a Dunnett’s multiple comparison test comparing values to the average of the water-treated samples.

**Figure 7:**
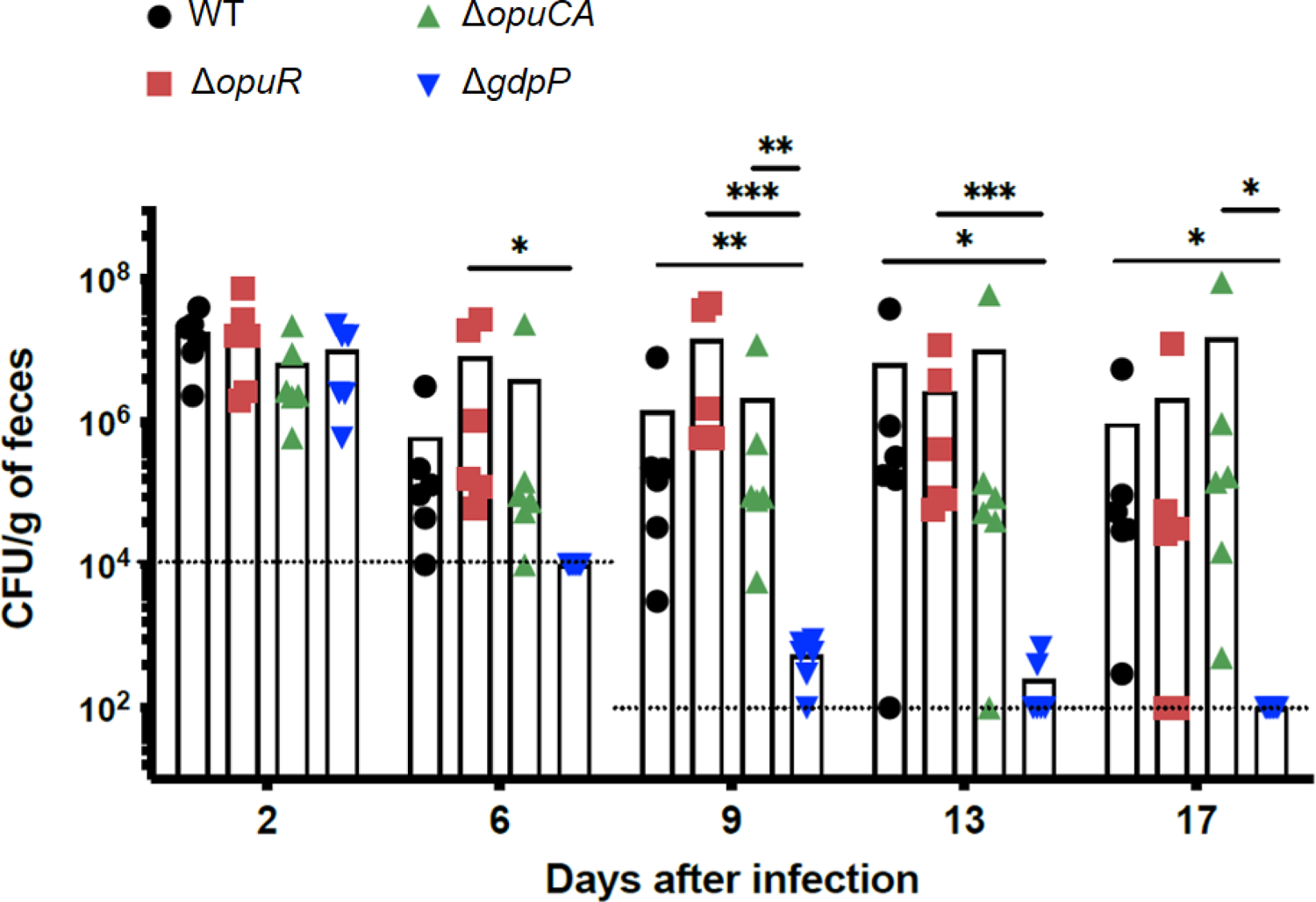
GdpP but not OpuR or OpuCA is required by *C. difficile* to sustain an infection of the murine gut. Mice were inoculated with 5×10^4^ spores of 630Δ*erm*, Δ*opuR*, Δ*opuCA* or Δ*gdpP*. Feces were collected on the indicated days and plated on BHI + 3% defibrinated horse blood + 0.1% taurocholate to assess total number of CFUs (vegetative cells + spores). *n*=6 mice; * *P* < 0.05; ** *P* < 0.01 and \*\*\**P* < 0.001 by a 0001 by a two-way ANOVA followed by a Dunnett’s multiple comparison test.

### GdpP but not OpuR or OpuCA is required for *C. difficile* persistence in the murine gut

Given the importance of GdpP, OpuR and OpuCA for resistance to osmotic and bile salts stresses, we hypothesized that they would impact initial colonization of the host and/or persistence in the gut. We therefore examined the impact of the corresponding mutants on the ability of *C. difficile* to colonize in an antibiotic-treated mouse model of CDI. Mice were pretreated with clindamycin and subsequently infected orally with 5×10^4^ spores of either 630Δ*erm,* Δ*gdpP*, Δ*opuR* or Δ*opuCA*. The intestinal burden of *C. difficile* was monitored by collecting feces from the mice over a 17-day period and enumerating colonies on selective medium. The number of total CFUs recovered from the feces of mice infected with the wild type or the mutant strains were all similar after 2 days of inoculation, suggesting that none of these genes is required to establish colonization of the gastrointestinal tract. At day 6, mice infected with the wild type strain showed a 1.5 log decrease in total CFUs compared to day 2 and then maintained this level of colonization throughout the time of the experiment. A similar pattern of colonization was observed for both the Δ*opuR* and Δ*opuCA* mutant strains. It is however worth noting that mice infected with Δ*opuR* had a higher bacterial burden than those infected with the wild type on day 9 after inoculation, although the differences did not reach statistical significance due to high variability among mice infected with the wild type strain. In contrast, mice infected with the Δ*gdpP* mutant showed a sharp decrease in CFU at days 6, 9 and 13 and completly cleared the bacteria at day 17. These data indicate that, under our experimental conditions, GdpP, unlike OpuR and OpuCA is essential for the persistence of *C. difficile* in the intestinal environment.

## DISCUSSION

In the past decade, the nucleotides secondary messenger molecule c-di-GMP has been found to regulate many important functions in *C. difficile*, including motility, adhesion, biofilm formation, and toxin expression (*59–66*). On the other hand, while *C. difficile* was previously shown to have an active DAC-encoding gene in its genome (*67*), production and roles of c-di-AMP had not been further investigated in this important enteropathogen prior to our study. Here, we demonstrated for the first time that *C. difficile* encodes 2 DACs and at least one PDE involved in synthesis and degradation of c-di-AMP, respectively. Decreased c-di-AMP concentrations caused by *cdaA* deletion were found to affect cell wall homeostasis resulting in increased susceptibility to peptidoglycan-targeting β-lactam antibacterial drugs. A direct connection between c-di-AMP synthesis and peptidoglycan biosynthesis has been established in *L. lactis*, *B. subtilis* and *S. aureus* (*68–71*). In these bacteria, the glucosamine-6-phosphate mutase GlmM, an enzyme for the production of the essential peptidoglycan synthesis intermediate glucosamine-1-P has been shown to form a complex with CdaA and to regulate CdaA activity (*72*). In numerous firmicutes, the CdaA-encoding gene is located in the same operon as *glmM*, together with *cdaR* coding for a cyclase regulator. Moreover, coexpresssion of the 3 genes from an upstream promoter has been shown in *S. aureus*, even though a second promoter was identified in front of *glmM* (*73*). In *C. difficile*, *glmM* is separated by a 7 genes operon (*CD0112-0118*) from the *cdaA-cdaR* gene cluster. Further studies are therefore needed to investigate the impact of a decreased expression of *cdaA-cdaR* on the transcription levels of *glmM* and vice-versa. We also found that elevated c-di-AMP concentrations promote biofilm formation in *C. difficile* as previously shown in several streptococci (*58, 74, 75*). In *S. mutans*, control of biofilm formation by c-di-AMP is mediated by the regulation of the expression of *gtfB*, encoding an enzyme responsible for the production of water-insoluble glucans. In contrast, accumulation of c-di-AMP in *B. subtilis* was shown to inhibit biofilm formation by affecting the activity of SinR (*56*). Factors connecting c-di-AMP and biofilm formation remain to be determined in *C. difficile*.

In this study, we demonstrated that the essential function of c-di-AMP in regulating osmotic homeostasis is conserved in *C. difficile*. The mechanism involves the binding of c-di-AMP to potassium transporter systems (Ktr and Kdp) and to the transcriptional regulator OpuR, which represses the expression of the *opuCAC* operon encoding a compatible solute transporter. OpuR contains a C-terminal TrkA_C domain (Pfam02080), which is also found in BusR, the Ktr family of K^+^ transporters and the K^+^ exporter CpaA and has been shown to be a c-di-AMP binding domain (*20, 21, 28, 38, 40, 42, 44, 76-79*). Deletion of *opuR* increased expression of the *opuCAC* operon, conferring a higher resistance to an hyperosmotic stress compared to the wild-type. Moreover, an overexpression of *opuCAC* had the same effect and a strain lacking the OpuCAC transporter was more susceptible than the wild type to NaCl. In line with our data, an osmotic shock with 1.5% NaCl for 1 h was found to consistently induce the expression of *opuCAC* and increase the abundance of the corresponding proteins in *C. difficile* strain 630 and ribotype 27 strains CD0196 (RefSeq_*opuCAC*: CD196_0850-0851) and QCD32g58 (RefSeq_*opuCAC*: CdifQ_04000997-04000998) (*80, 81*). Based on our results, deletion of *opuR* results in an uncontrolled compatible solute influx through the constant expression of *opuCAC* in *C. difficile* and this leads to an elevation of the c-di-AMP concentrations. The same observation was reported in *L. lactis* (*21*). In contrast, c-di-AMP concentrations are decreased in an *opuCA* mutant. These data thus suggest that sensing of fluctuations of the cellular turgor constitutes a signal to adjust the intracellular concentration of c-di-AMP in *C. difficile*.

A new and important finding from our study is that bile salt exposure rapidly stimulates c-di-AMP degradation and we clearly demonstrated that the OpuCAC-mediated osmolyte uptake plays an important role in bile salt tolerance. The mechanism involved seems to be identical to the one conferring resistance to NaCl. In response to a hyperosmotic or a bile salt stress, the cells lower their c-di-AMP concentrations, which triggers a relief of *opuCAC* genes repression by OpuR and an increased uptake of protective compatible solutes. Consistent with the role of OpuCAC in bile salt resistance, expression of *opuCA* was induced 12-fold in biofilms obtained after 48 h of growth in the presence of deoxycholate compared to the absence of bile salts (*82*). In addition, the abundance of OpuCA and OpuCC proteins was increased in *C. difficile* 630Δ*erm* after 90 min exposure to either the primary bile salts cholate or chenodeoxycholate, or the secondary bile salts deoxycholate or litocholate (*83*). In *L. monocytogenes*, the carnitine transporter OpuC is also involved in bile tolerance and *opuC* genes transcription is induced in response to bile salts (*61*). C-di-AMP was shown to bind the cystathionine beta-synthase (CBS) domain of OpuCA in *L. monocytogenes* but no link between c-di-AMP and bile tolerance has yet been established in this bacterium to our knowledge (*46*).

Expression of *opuC* genes is regulated by the stress-inducible sigma factor σ^B^ in *L. monocytogenes*, with putative σ^B^ promoter motifs identified upstream *opuCA* (*84, 85*). The osmotic induction of *opuC* genes has been shown to be strongly σ^B^ dependent and tolerance to bile salts is also mediated by σ^B^ in *L. monocytogenes* (*86, 87*). In *C. difficile*, the *opuCAC* operon is also positively controlled by σ^B^ and a σ^B^-dependent promoter was identified upstream *opuCA* (*88, 89*). However, a strain lacking this Sigma factor showed no growth defect compared to the wild type in the presence of NaCl or bile salts (*89*). A putative σ^A^ promoter but no σ^B^ promoter region was identified upstream *opuR* of *C. difficile* (*90*). It is therefore conceivable that OpuR still modulates *opuCAC* expression in response to fluctuations in the c-di-AMP concentrations in the absence of σ^B^. The c-di-AMP signaling pathway would thus provide a specific response to osmotic and bile salt stresses independently of the activation of the general stress response by σ^B^ in *C. difficile*.

Due to their amphipathic nature, bile salts are antibacterial compounds that act as natural detergents and disrupt bacterial membranes (*91, 92*). In *Enterococccus faecalis* and *Propionibacterium freudenreichii*, a pretreatment with a sublethal dose of bile salts or of the detergent SDS conferred a similar protection against lethal levels of bile salts, demonstrating that the physiological responses to bile salts and SDS are closely related (*93, 94*). We showed in this study that OpuR and OpuCA also play an important role in SDS and Triton X-100 tolerance suggesting that intracellular c-di-AMP concentrations are decreased in response to the detergent activity of bile salts to confer protection. Supporting our data, c-di-AMP-specific PDE genes mutants with elevated c-di-AMP concentrations were more sensitive to Triton X-100 than their respective wild type in *Bacillus anthracis* and *Streptococccus mitis* (*95, 96*).

In *L. monocytogenes*, cell adaptation to either SDS or to NaCl conferred a similar high cross-protection against lethal levels of bile salts (*97*). Likewise, a pre-treatment of *E. faecalis* cells with subinhibitory concentrations of NaCl induced tolerance against lethal levels of SDS or bile salts challenges (*98*). Both osmotic and detergent stresses alter membrane characteristics, the former by modifying the cellular turgor (*99*). This leads us to hypothesize that the molecular mechanisms by which the c-di-AMP concentrations are modulated in response to osmotic and bile salt stresses sensing are likely the same and associated to the maintenance of membrane integrity. As previously discussed by Pham *et al* (*21*), an attractive possibility to link membrane alterations to the fluctuations of c-di-AMP concentrations would be that changes in membrane characteristics are directly sensed by the membrane-bound enzymes CdaA and GdpP through their hydrophobic domains.

The colonization and virulence of several c-di-AMP-producing pathogens are affected by altering c-di-AMP homeostasis (*30, 31, 36, 58, 74, 95, 100, 101*). Likewise, *C. difficile* lacking *gdpP* was deficient in long-term colonization of the mice intestine, presumably because of the exacerbated susceptibility of this strain to environmental stresses. In contrast, OpuC was shown to not be required for *C. difficile* maintenance in the gut in our experimental conditions. This difference is probably due to the fact that c-di-AMP induces osmotic stress adaptation via the transport of K^+^ and compatible solute transporters, and bile salt resistance exclusively through the control of *opuC* expression. Thus, these data suggest that the gut persistence defect of Δ*gdpP* is caused by its inability to adapt to a high osmotic pressure rather than the presence of bile salts. However, we cannot exclude that susceptibility to other stresses unexplored in this study or that an indirect effect of the *gdpP* deletion on the cell physiology contributes to this phenotype. It is now recognized that the biotransformation of primary bile acids produced by the liver into secondary bile acids by members of the gut microbiota plays an important role in the mechanism of colonization resistance (*102–105*). Before antibacterial drugs treatment, secondary bile salts inhibit *C difficile* growth. However, the gut microbiota is altered by an antibiotic treatment leading to an increased concentration of the primary bile salts, which supports *C difficile* spore germination and outgrowth (*7, 102, 106, 107*). In our study, mice were treated with clindamycin to make them susceptible to *C. difficile* 630Δ*erm* infection which will decreases gut bacteria diversity and affect bile salt profile (*108*). Thus, our experimental conditions cannot evaluate the full impact of the *opuCA* deletion on the colonization and persistence of *C. difficile* given the lack of secondary bile salts. Additional work with a different animal model is required to determine the contribution of the OpuC transporter and OpuR regulator to *C. difficile* gut persistence in the presence of secondary bile salts. Nevertheless, our work identified c-di-AMP as a crucial signaling molecule regulating adaptation to the host environment, colonization and, for the first time, tolerance to bile salts.

## MATERIALS AND METHODS

### Bacterial strains and culture conditions

*C. difficile* and *E. coli* strains used in this study are presented in Table S1. *E. coli* strains were grown in LB broth, and when needed, with ampicillin (100 μg/ml) or chloramphenicol (15 μg/ml). *C. difficile* strains were grown in an anaerobic chamber (Jacomex) under an anaerobic atmosphere (5 % H_2_, 5 % CO_2_, and 90 % N_2_) in TY (*109*), *C. difficile* minimal medium (CDMM) (*110*) or Brain Heart Infusion (BHI, Difco) media. When necessary, cefoxitin (Cfx; 25 μg/ml), cycloserine (Cs; 250 μg/ml) and thiamphenicol (Tm; 7.5 μg/ml) were added to *C. difficile* cultures. A potassium-free medium derived from CDMM was used to study potassium requirements of *C. difficile* strains (Table S2). Potassium chloride was added as indicated. The non-antibiotic analog anhydrotetracycline (ATc, Sigma-Aldrich) was used for induction of the *P_tet_* promoter of pRPF185 vector derivatives in *C. difficile* (*111*). Growth curves were obtained in 96 wells microplates at 37°C and automatic recording of OD_600_ every 30 minutes using a plate reader (Promega GloMax Explorer).

### Plasmid and strain construction

All primers used in this study are listed in Table S3. For deletions, allele exchange cassettes were designed to have between 800 and 1050 bp of homology to the chromosomal sequence in both up- and downstream locations of the target gene. The homology arms were amplified by PCR from *C. difficile* strain 630 genomic DNA and purified PCR products were directly cloned into the PmeI site of pMSR vector using NEBuilder HiFi DNA Assembly (New England Biolabs). All pMSR-derived plasmids were initially transformed into *E. coli* strain NEB10β (New England Biolabs), and sequences of all inserts were verified by sequencing. Plasmids were then transformed into *E. coli* HB101(RP4) and transferred by conjugation into the appropriate *C. difficile* strains. Transconjugants were selected on BHI supplemented with cycloserine, cefoxitin, and thiamphenicol. Allelic exchange was performed following the procedure previously published (*112*). For the expression of recombinant KtrA, KdpD and OpuR, DNA sequence of the corresponding genes was amplified by PCR from *C. difficile* strain 630 genomic DNA and cloned into *BamHI* and *PstI* sites of pQE30 expression vector (Qiagen). All pQE30-derived plasmids were initially transformed into *E. coli* strain NEB10β (New England Biolabs), and sequences of all inserts were verified by sequencing. Plasmids were then transformed into *E. coli* strain XL1-blue for protein expression. For inducible expression of *cdaA* and *gdpP* in *C. difficile*, DNA sequence of the corresponding genes and their preceding RBS was amplified by PCR from *C. difficile* strain 630 genomic DNA and cloned into *XhoI* and *BamHI* sites of pDIA6103 vector. For constitutive expression of *opuCAC* in *C. difficile*, the *P_tet_* promoter and regulatory region were excised from pDIA6103 by inverse PCR. The *P_cwp2_* promoter and the *opuCAC* coding sequence with its preceding RBS were amplified by PCR from *C. difficile* strain 630 genomic DNA and cloned into the modified pDIA6103 using NEBuilder Hifi DNA Assembly (New England Biolabs). The resulting pDIA6103 derivative plasmid was transformed into the *E. coli* HB101 (RP4) and subsequently mated with *C. difficile* 630Δ*erm* strain. Transconjugants were selected on BHIS supplemented with cycloserine, cefoxitin, and thiamphenicol.

### RNA isolation and quantitative reverse-transcriptase PCR

Total RNAs were isolated from *C. difficile* strains after 4 h of growth in TY medium. Total RNA extraction, cDNA synthesis and real-time quantitative PCR were performed as previously described (*113*). In each sample, the quantity of cDNAs of a gene was normalized to the quantity of cDNAs of the *dnaF* gene (*CD1305*) encoding DNA polymerase III. The relative change in gene expression was recorded as the ratio of normalized target concentrations (threshold cycle [ΔΔ*C_T_*] method) (*114*).

### RNA sequencing

Transcriptomic analysis for each condition was performed using 4 independent total RNA preparations using methods described before (*82*). Briefly, the RNA samples were first treated using Epicenter Bacterial Ribo-Zero kit. This depleted rRNA fraction was used to construct cDNA libraries using TruSeq Stranded Total RNA sample prep kit (Illumina). Libraries were then sequenced by Illumina NextSeq 500. Cleaned sequenced reads were aligned to the reannotated *C. difficile* strain 630 (*58*) for the mapping of the sequences using Bowtie (Version 2.1.0). DEseq2 (version 1.8.3) was used to perform normalization and differential analysis using values of the Δ*gdpP* strain as a reference for reporting the expression data of the Δ*gdpP*Δ*opuR* strain. Genes were considered differentially expressed if they had ≥ 2-fold increase or decrease in expression and an adjusted *p*-value (*q* value) ≤0.05.

### Phase-contrast microscopy

Bacterial cells were observed at 100x magnification on an Axioskop Zeiss Light Microscope. Cell length of 100 cells was measured for each strain using ImageJ software (*115*).

### Antibiotic susceptibility tests

Susceptibility tests for antibacterial drugs were conducted using disk diffusion assay. Overnight cultures of *C. difficile* strains were diluted to a starting OD_600_ of 0.05, grown to an OD_600_ of 1 and 100 μl of the culture was spread on a TY agar plate. A disk of 10 μg imipenem, 30 μg cefepime or 30 μg moxalactam (Bio-Rad) was then placed on top of the plate. The zone of growth inhibition was measured after incubation for 24 h at 37°C.

### Biofilm formation

To generate biofilms, overnight cultures of *C. difficile* were diluted 1:100 into fresh BHISG (BHI supplemented with 0.1% L-cysteine, 5 mg/ml yeast extract and 100 mM glucose), 1 mL per well was deposited in 24-well polystyrene tissue culture-treated plates (Costar, USA) and the plates were incubated at 37 °C in anaerobic environment for 24 h. Biofilm biomass was measured using a crystal violet as described elsewhere (82). Briefly, spent media is removed by inversion, and wells are washed twice with PBS, dried, stained with crystal violet for 2 min and wash twice with PBS. Crystal violet was the solubilized with a 75% ethanol solution and the absorbance was measured at a λ_600nm_ with a plate reader (Promega GloMax Explorer, Promega, France).

### Measurement of intracellular c-di-AMP concentrations

For quantification of the c-di-AMP concentrations in the mutant strains, overnight cultures of *C. difficile* strains were diluted in TY to a starting OD_600_ of 0.05 and grown for 4 h at 37 °C. A 10-ml culture aliquot was harvested by centrifugation at 5000 × *g* for 10 min at 4 °C. Nucleotides were extracted from the pellet with methanol/acetonitrile/milli-Q water (40:40:20) and the c-di-AMP detected and quantified by LC-MS/MS, as described previously (*116*). For normalization purposes, a 1-ml aliquot from the same culture was taken and pelleted for 10 min at 8,000 x*g* at 4 °C. Bacterial pellets were frozen and then thawed, resuspended in 1 ml PBS and incubated at 37 °C for 45 min to lyse the cells. The samples were centrifuged for 5 min at 10,000 x*g* and the protein content of the supernatant was determined using a BCAassay kit (Thermo Scientific). The c-di-AMP concentrations are presented as ng c-di-AMP/mg *C. difficile* protein.

The previously described energized cell suspension method was used to determine the impact of an osmotic or a bile salt stress on the c-di-AMP concentrations (*21*). A 25 ml culture of *C. difficile* 630Δ*erm* grown in TY until OD_600_ of 0.7 was harvested by centrifugation at 5,000 × *g* for 10 min and washed twice with low osmolarity 1/10 KPM buffer (0.01M K_2_HPO_4_ adjusted to pH 6.5 with H_3_PO_4_ and 1mM MgSO_4_.7H_2_O). Cells were then resuspended in 3.75 ml 1/10 KPM and 20 mM D-glucose was added to the cell suspension before incubating for 10 min at 37°C to stimulate ATP production required for c-di-AMP synthesis. The cell suspension was then split into 750 µL aliquots. An aliquot was collected to represent t_0_ and 50 µL of either water, NaCl (300 mM final), deoxycholate (0.028% final) or cholate (0.4% final) was added to the other aliquots before an additional 10 min incubation at 37°C. When the desired time point was reached, samples were mechanically lysed using beads and a FastPrep 5G system (MP Biomedicals) and the supernatant was collected after centrifugation (5 min at 16,000 × g). A small aliquot was removed, to measure protein concentration using a Pierce BCA protein assay kit (Thermo Scientific). The remainder of the sample was heated to 95°C for 10 min and used to quantify c-di-AMP using a c-di-AMP ELISA kit (Cayman Chemical) according to the recommendations of the supplier. The c-di-AMP concentrations are presented as ng c-di-AMP/mg *C. difficile* protein.

### Differential Radial Capillary Action of Ligand Assay (DRaCALA)

Interaction between c-di-AMP and target proteins was tested by DRaCALA on whole *E*. *coli* protein extract (*20*). Expression of the KtrA, KdpD and OpuR tagged proteins was induced with 1 mM IPTG for 6 h at 30°C. Bacterial pellets from 1 ml culture were resuspended in 100 μl binding buffer (40 mM Tris pH 7.5, 100 mM NaCl, 10 mM MgCl2, 0.5 mg/ml lysozyme, 20 μg/ml DNase) and lysed by 3 freeze-thaw cycles. For DRaCALA, 1 nM [^32^P]-c-di-AMP, synthetized as previously described (*20*), was added to the whole protein extract, incubated at room temperature for 5 min, and 2.5 μl was spotted onto nitrocellulose membrane. For competition assays, reactions were incubated with 150 μM of nonlabeled nucleotides (c-di-AMP, c-di-GMP, cAMP, cGMP, AMP, and ATP; BioLog Life Science Institute, Germany) for 5 min at room temperature prior to addition of 1 nM [^32^P]-c-di-AMP. Samples were spotted on nitrocellulose after 5 min reaction at room temperature. Radioactive signal was detected with a Typhoon system (Amersham).

### Conventional Mouse Infection Studies

*C. difficile* spore inoculums were generated by plating 200 µL of an overnight culture of *C. difficile* grown on SMC medium (9% Bacto peptone, 0.5% proteose peptone, 0.15% Tris base, 0.1% ammonium sulphate) in SMC agar and incubating them at 37°C for 7 days in an anaerobic chamber. Spores were then harvested in 2 ml of ice-cold sterile water and purified by centrifugation using a HistoDenz (Sigma-Aldrich) gradient (*117*).

6-week old conventional C57BL mice (Janvier Labs) were singly housed and acclimated for a week prior to treatment with clindamycin (10mg/kg; Sigma Aldrich)) via intraperitoneal (IP) injection. 24 hours post-clindamycin treatment, groups of 6 mice were challenged with 5×10^4^ wild-type or mutant *C. difficile* spores via oral gavage. To assess bacterial persistence, fecal pellets were collected over a 17 days period (days 2, 6, 9, 13 and 17). Fecal pellets were homogenized in the anaerobic hood in 1x pre-reduced PBS at a concentration of 100 mg/ml, serially diluted and plated on BHI agar containing 3% defibrinated horse blood and 0.1% taurocholate to assess total number of CFUs.

## Supplementary Materials

Fig. S1. C-di-AMP is involved in osmotolerance.

Fig. S2. C-di-AMP and K^+^ uptake in *C. difficile*.

Fig. S3. OpuR represses *opuCA* expression.

Fig. S4. C-di-AMP modulates resistance to the detergent action of bile salts by controlling the OpuCAC-mediated import of osmolytes through OpuR.

Table S1. Strains and plasmids used in this study.

Table S2. Potassium-free medium derived from CDMM.

Table S3. Oligonucleotides used in this study.

## ACKNOWLEDGMENTS

We thank Lena Winat and Flor Saporta for technical support and Isabelle Martin-Verstraete, Nicolas Kint and Marc Monot for helpful discussions. We would also like to thank Pierre-Alexandre Kaminski for the gift of the purified diadenylate cyclase DisA from *B. subtilis* and for helpful discussions.

## Author contributions

The following author contributions were made.

Conceptualization: BD, JP

Methodology: BD, JP

Investigation: MO, AH, PBS, PAS, YDNT,ND, RS, JP

Visualization: MO, AH, RS, OS, LB, BD, JP Funding acquisition: JP, BD, OS

Project administration: JP, BD Supervision: JP, BD

Writing – original draft: JP

Writing – review & editing: MO, AH, PBS, PAS, YDNT, RS, OS, BD, JP

## Data and materials availability

All data are available in the main text or the supplementary materials.

