## Supplemental materials for "c-di-AMP signaling is required for bile salts resistance and long-term colonization by *Clostridioides difficile*"

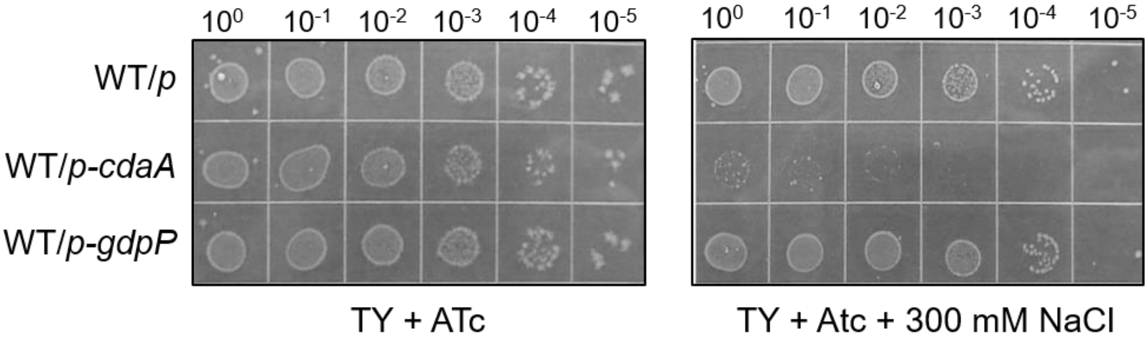


**Figure S1: C-di-AMP is involved in osmotolerance.** Growth of *C. difficile* 630Δ*erm* carrying an empty vector (WT/p) or a vector expressing *cdaA* (WT/*p-cdaA*) or *gdpP* (WT/*p-gdpP*) under the control of the $P_{tet}$ promoter on TY agar or TY agar + 300 mM NaCl in presence of 250 ng/ml ATc after 24 h incubation at 37°C. Spots (5µl) are from cells grown overnight in TY and serially diluted. Data are representative of 3 experiments.

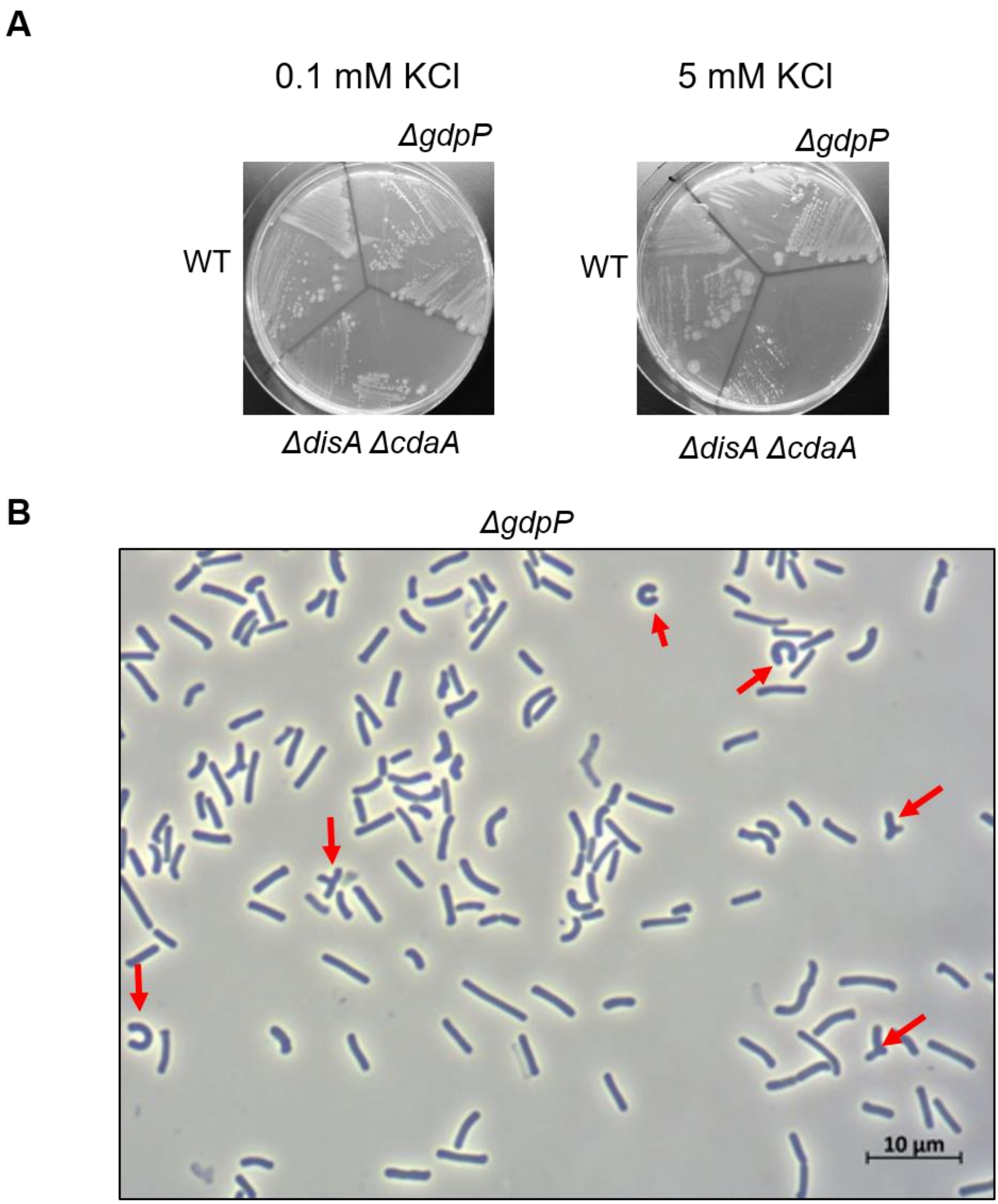


**Figure S2: C-di-AMP and $K^+$ uptake in *C. difficile*. (A)** Growth of *C. difficile* wild type, Δ*gdpP* and Δ*disA*Δ*cdaA* strains on CDMM agar plates containing 0.1 mM or 5 mM KCl after 48 h at 37 °C. Data are representative of 3 experiments. (B) Phase contrast microscopy image of *C. difficile* Δ*gdpP* grown for 48 h at 37 °C on CDMM agar plates containing 1 µM KCl. Data are representative of 3 experiments.

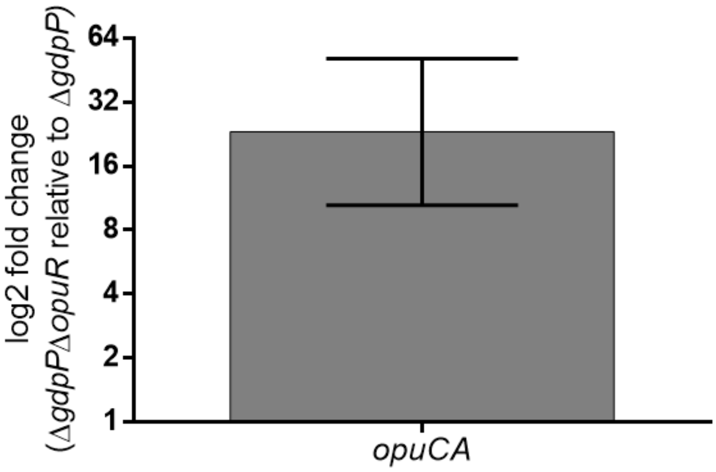


**Figure S3: OpuR represses *opuCA* expression.** Comparative qRT-PCR analysis of *opuCA* transcript levels in Δ*gdpP* and Δ*gdpP* Δ*opuR* grown in TY medium for 4 h. Means and SEM are shown; *n* = 4.

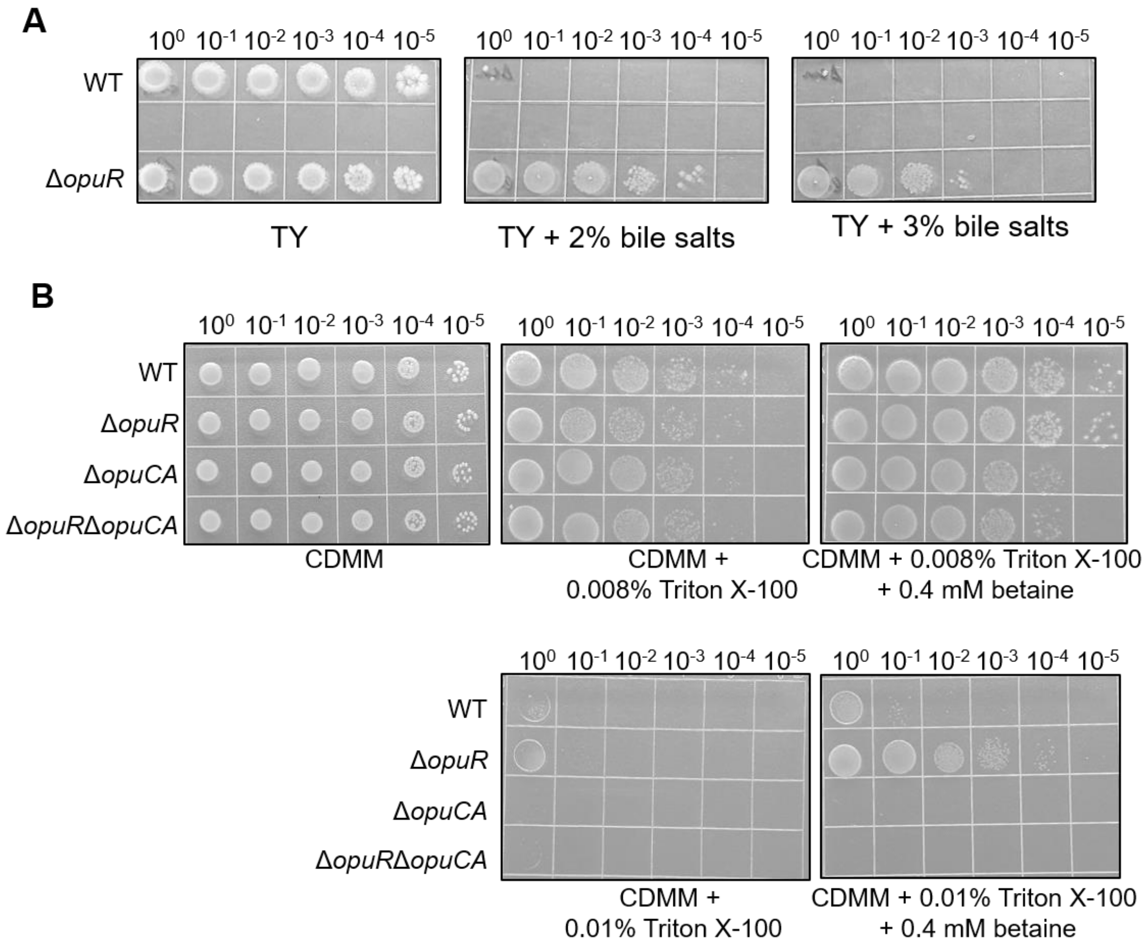


**Figure S4: C-di-AMP modulates resistance to the detergent action of bile salts by controlling the OpuCAC-mediated import of osmolytes through OpuR. (A)** Growth of *C. difficile* wild type and mutant strains on TY agar, TY agar + 2% bile salts or TY agar + 2% bile salts after 48 h incubation at 37°C. Spots (5µl) are from cells grown overnight in TY and serially diluted. Data are representative of 3 experiments. **(B)** Growth of *C. difficile* wild type and mutant strains on CDMM agar, CDMM agar + 0.008% Triton X-100, CDMM agar + 0.008% Triton X-100 + 0.4 mM glycine betaine, CDMM agar + 0.01% Triton X-100, CDMM agar + 0.01% Triton X-100 + 0.4 mM glycine betaine after 48 h incubation at 37°C. Spots (5µl) are from log-phase cells grown in TY, washed once in 0.9% saline and serially diluted. Data are representative of 3 experiments.

**Table S1. Strains and plasmids used in this study.**

| Strain | Genotype | Origin |
|---|---|---|
| ***E. coli*** | | |
| NEB-10 beta | *Δ(ara-leu) 7697 araD139 fhuA ΔlacX74 galK16 galE15 e14- ϕ80*d*lacZΔM15 recA1 relA1 endA1 nupG rpsL* (Str$^R$) *rph spoT1 Δ(mrr-hsdRMS-mcrBC)* | New England Biolabs |
| HB101 (RP4) | *supE44 aa14 galK2 lacY1* Δ(*gpt-proA*) 62 *rpsL20 (*Str$^R$*)xyl-5 mtl-1 recA13* Δ(*mcrC-mrr*) *hsdS*$_B$ ($r_B$-$m_B$-) RP4 (Tra$^+$ IncP Ap$^R$ Km$^R$ Tc$^R$) | Laboratory stock |
| XL1-blue | Δ(*mcrA*)*183* Δ(*mcrCB-hsdSMR-mrr*)*173 endA1 supE44 thi-1 recA1* gyrA96 relA1 lac *[F′* proAB lacI$^q$Z*Δ*M15 *Tn*10 *(Tet$^r$)]* | Laboratory stock |
| EC1047 | XL1-blue strain carrying pDIA6675 for expression of 6xHis-tagged OpuR | This work |
| EC1046 | XL1-blue strain carrying pDIA6676 for expression of 6xHis-tagged KdpD | This work |
| EC1045 | XL1-blue strain carrying pDIA6759 for expression of 6xHis-tagged KtrA | This work |
| ***C. difficile*** | | |
| 630Δ*erm* | 630 Δ*ermB* | Laboratory stock (*1*) |
| CDIP369 (630/p) | 630 Δ*erm* strain carrying pDIA6103 | Laboratory stock |
| CDIP755 | 630 Δ*erm* strain carrying pDIA6504 for inducible expression of *cdaA* | This work |
| CDIP758 | 630 Δ*erm* strain carrying pDIA6512 for inducible expression of *gdpP* | This work |
| CDIP660 | 630 Δ*erm* Δ*cdaA* | This work |
| CDIP772 | 630 Δ*erm* Δ*disA* | This work |
| CDIP773 | 630 Δ*erm* Δ*gdpP* | This work |
| CDIP1199 | 630 Δ*erm* Δ*disA* Δ*cdaA* | This work |
| CDIP1291 | 630 Δ*erm* Δ*opuR* | This work |
| CDIP1305 | 630 Δ*erm* Δ*gdpP* Δ*opuR* | This work |
| CDIP1362 | 630 Δ*erm* Δ*opuCA* | This work |
| CDIP1364 | 630 Δ*erm* Δ*opuR* Δ*opuCA* | This work |
| **Plasmid** | | |
| pQE-30 | For expression of 6xHis-tagged proteins | Qiagen |
| pRPF185 | $P_{tet}$-*gusA* Tm$^{R,}$ expression and cloning vector | (*2*) |

| | | |
|---|---|---|
| pMSR | Allele exchange in *C. difficile* 630 | This work |
| pDIA6103 | pRPF185 Δ*gus* vector derivative | (*3*) |
| pDIA6504 | pDIA6103 derivative for inducible *cdaA* expression | This work |
| pDIA6512 | pDIA6103 derivative for inducible *gdpP* expression | This work |
| pDIA6466 | pMSR derivative for *cdaA* deletion | This work |
| pDIA6510 | pMSR derivative for *gdpP* deletion | This work |
| pDIA6511 | pMSR derivative for *disA* deletion | This work |
| pDIA6675 | pQE-30 derivative for *busR* expression | This work |
| pDIA6676 | pQE-30 derivative for *kdpD* expression | This work |
| pDIA6759 | pQE-30 derivative for *ktrA* expression | This work |
| pDIA6854 | pMSR derivative for *disA* deletion | This work |

**Table S2. Potassium-free medium derived from CDMM.**

| Stock solution component | Concn in stock solution (mg/ml) | Final concn in CDMM (mg/ml) |
|---|---|---|
| **Amino acids (5X)** | | |
| Histidine | 0,5 | 0,1 |
| Tryptophane | 0,5 | 0,1 |
| Glycine | 0,5 | 0,1 |
| Arginine | 0,5 | 0,1 |
| Methionine | 0,5 | 0,1 |
| Threonine | 0,5 | 0,1 |
| Valine | 0,5 | 0,1 |
| Isoleucine | 0,5 | 0,1 |
| Leucine | 5 | 1 |
| Cysteine | 2.5 | 0.5 |
| Proline | 4 | 0,8 |
| **Salts** | | |
| $Na_2HPO_4$ | 50 | 0.9 |
| **Glucose (20X)** | | |
| D-Glucose | 200 | 10 |
| **Trace salts (50X)** | | |
| $(NH_4)_2SO_4$ | 2.0 | 0.04 |
| CaCl2 2H2O | 1.3 | 0.026 |
| MgCl2 6H2O | 1.0 | 0.02 |
| MnCl2 4H2O | 0.5 | 0.01 |
| CoCl2 6H2O | 0.05 | 0.001 |
| **Iron (100X)** | | |
| $FeSO_4$ $7H_2O$ | 0.4 | 0.004 |
| **Vitamins** | | |
| D-Biotin (100000X) | 1 | 1x10-5 |
| Calcium-D-panthothenate (1000X) | 1 | 1x10-3 |
| Pyridoxine (10000X) | 1 | 1x10-4 |

**Table S3. Oligonucleotides used in this study.**

| Name | Sequence (5'-3')[1] | Description |
|---|---|---|
| **pDIA6103 cloning** | | |
| IMV507 | GGGATTTCTCACATAAAATAGAG | 5'pDIA6103 insert screening |
| IMV508 | TAAAATAAGCTTGATCGTAGCG | 3'pDIA6103 insert screening |
| JP248 | AGCACTCGAGGTTATCAAACAGGAGGATAAAAC | 5'*cdaA-XhoI* |
| JP249 | AGCAGGATCCTCATTTAAATATACCACCTTTGAA | 3'*cdaA-BamHI* |
| JP252 | AGCACTCGAGGCTATAAATCCACTTAGGAGGATAG | 5'*gdpP-XhoI* |
| JP253 | AGCAGGATCCCTATTCTTCCTCCTCCAAATAC | 3'*gdpP-BamHI* |
| JP721 | CTTCTCGAGAGGATCCTATAAG | 5' $P_{tet}$ and its regulatory element removal from pDIA6103 by inverse PCR |
| JP722 | GTGAAAGTGGGTCTTAAGGTAC | 3' $P_{tet}$ and its regulatory element removal from pDIA6103 by inverse PCR |
| JP723 | accttaagacccactttcacCTGGATTATTAAAAAAAATTACAATTTTG | 5' $P_{cwp2}$ |
| JP716 | taaaaatttgGAAAAAGGGGATGATAAATTATG | 3' $P_{cwp2}$ |
| JP717 | cccctttttcCAAATTTTTATTATTTTTTCCTTATTTACC | 5' *opuCAC* |
| JP724 | tataggatcctctcgagaagGAAATGATAAAATATATTTGATTTCATATAAAAAAG | 3' *opuCAC* |
| **pQE-30 cloning** | | |

| | | |
|---|---|---|
| JP386 | AGCAGGATCCAAGCAATATATAGTCATTGGGTG | *5'ktrA-BamHI* |
| JP387 | AGCACTGCAGTTATCTTCTTCTAATTATCCCCTTATT | *3'ktrA-PstI* |
| JP388 | AGCAGGATCCGATAAAAACTATACGATGCCAATATAC | *5'opuR-BamHI* |
| JP389 | AGCACTGCAGTTATTTTTCGTTTTCATATAAAAACAT | *3'opuR-PstI* |
| JP390 | AGCAGGATCCGATAGACCTAATCCTGATATTCTATTAG | *5'kdpD-BamHI* |
| JP391 | AGCACTGCAGCTATTCATTTTCCTTAGGCAAA | *3'kdpD-PstI* |
| **pMSR cloning** | | |
| JP449 | CGTTTTGTAAACGAATTGC | 5'pMSR insert screening |
| JP450 | CTCACGTTAAGGGATTTTG | 3'pMSR insert screening |
| JP053 | cggtgtttttgttaccctaagtttGTAGTTCAGTTGGTTAGAATG | 5' left arm Δ*cdaA* |
| JP131 | taaatataccTCCATTTAAAAAAGAATTTATATCTAACAC | 3' left arm Δ*cdaA* |
| JP132 | tttaaatggaGGTATATTTAAATGATAAATAGACTAAAAAATAATAC | 5' right arm Δ*cdaA* |
| JP133 | gattatcaaaaaggagtttGTCGCATCTTGTAACAAC | 3' right arm Δ*cdaA* |
| JP063 | GAAAAGTACAGCGTCTCTACATC | 5' Δ*cdaA* screening |
| JP138 | CGCTTTTTAGATTTAACAAATATCTC | 3' Δ*cdaA* screening |
| JP227 | ttttttgttaccctaagtttGAAGAAGAGCGGTTTACAAC | 5' left arm Δ*disA* |
| JP228 | tgtgtctatcCTCCATATTTATTGATTCCTCC | 3' left arm Δ*disA* |
| JP229 | aaatatggagGATAGACACATATGATGAATTTTAAATG | 5' right arm Δ*disA* |
| JP230 | agattatcaaaaaggagtttGAACAGAAGTAACGAGTACTTTTATAG | 3' right arm Δ*disA* |
| JP231 | CCAGTCAGTATAAACTCAATAGAGAG | 5' Δ*disA* screening |
| JP232 | CTTCACCCCCTAGTTATACTTATC | 3' Δ*disA* screening |
| JP234 | ttttttgttaccctaagtttGCAAGTGTTAGTGATATTACAG | 5' left arm Δ*gdpP* |
| JP235 | ataattatataaATGACTGCTTATAACAATATAGC | 3' left arm Δ*gdpP* |

| | | |
|---|---|---|
| JP236 | aagcagtcatTTATATAATTATTTGGACTAACATATAGTATC | 5’ right arm Δ*gdpP* |
| JP237 | agattatcaaaaaggagtttGGGGATTATCAGCAATGAAC | 3’ right arm Δ*gdpP* |
| JP238 | GATAGTTATGGATTCTCATGGTG | 5’ Δ*gdpP* screening |
| JP239 | CTTGTGTCTAACCTTTGCATC | 3’ Δ*gdpP* screening |
| JP485 | tttttgttaccctaagtttCCCCATAGCTAACTCTTTATC | 5’ left arm Δ*opuR* |
| JP486 | tttcgttttcGTATATTGGCATCGTATAGTTTTTATC | 3’ left arm Δ*opuR* |
| JP487 | gccaatatacGAAAACGAAAAATAAATTATATAATTTAGTG | 5’ right arm Δ*opuR* |
| JP488 | agattatcaaaaaggagtttGCCATTAGTAAAACCTTCTAG | 3’ right arm Δ*opuR* |
| JP489 | GCAAGACAGAATCATTTTTCTTCATTC | 5’ Δ*opuR* screening |
| JP491 | GAGAATTGAACATACCAATAAGC | 3’ Δ*opuR* screening |
| JP551 | tttttgttaccctaagtttGAGATAGCAAAGCAAATAAAAAAAG | 5’ left arm Δ*opuCA* |
| JP552 | agccaagaacCTCTATCATAATTTATCATCCCC | 3’ left arm Δ*opuCA* |
| JP553 | tatgatagagGTTCTTGGCTAATTTTTTTAACTTTATAC | 5’ right arm Δ*opuCA* |
| JP554 | agattatcaaaaaggagtttCCATAGTATCTTTTACAACATTATATG | 3’ right arm Δ*opuCA* |
| JP555 | GTAGCAATATTCAGACAAATAGTGAC | 5’ Δ*opuCA* screening |
| JP556 | CAGTTTCAGGCATCATGG | 3’ Δ*opuCA* screening |
| **quantitative PCR** | | |
| RT_polIII-F_Cdiff | TCCATCTATTGCAGGGTGGT | 5' *CD1305* |
| RT_polIII-R_Cdiff | CCCAACTCTTCGCTAAGCAC | 3' *CD1305* |
| IMV786 | TGTCAAGTGAGCCTTTGCAT | 5’ *CD0900 (opuCA)* |
| IMV787 | TCCAACCGAATTACGATTCA | 3’ *CD0900 (opuCA)* |

[1]Underlined bases indicate engineered restriction sites; lowercase bases indicate overlapping sequences
